# Functional divergence of Capicua isoforms explains differential tissue vulnerability in neurological disease

**DOI:** 10.1101/2025.11.05.686788

**Authors:** Hamin Lee, Esmeralda Villavicencio Gonzalez, Elias M. Rivera, Mark A. Durham, Ronald Richman, Elizabeth H.-Y. Chu, Kailey Xia, Hu Chen, Zhandong Liu, Surabi Veeraragavan, Binoy Shivanna, Huda Y. Zoghbi

## Abstract

Many neurological diseases impact specific brain regions despite widespread expression of the disease-related protein. Spinocerebellar ataxia type 1 (SCA1) primarily affects the cerebellum, though Ataxin-1 (ATXN1) is widely expressed. We previously showed that intensified interaction between mutant ATXN1 and Capicua (CIC) drives SCA1 pathogenesis in the cerebellum, whereas ATXN1 loss augments amyloid beta production in the hippocampus and cortex. CIC, however, forms a complex with ATXN1 and its paralog, Ataxin-1-like (ATXN1L)—yet knockout of either yields completely different phenotypes. To determine whether this could be due to CIC having two isoforms, we generated mice bearing either the long (CIC-L) or short (CIC-S) isoform. Loss of CIC-L led to cognitive deficits, whereas loss of CIC-S caused perinatal lethality, phenocopying, ATXN1 and ATXN1L knockout mice, respectively. Furthermore, CIC-L preferentially interacts with ATXN1, and CIC-S with ATXN1L. Our data underscore the importance of isoform–paralog interplay in studying regional vulnerability in neurodegenerative diseases.

## INTRODUCTION

One of the puzzling features of many neurological diseases is that they seem to affect particular brain regions or cell types even though the disease-related protein is expressed widely throughout the brain and often in other tissues as well (Fu et al. 2018; Kampmann 2024). Typically, the disease ramifies outward from that locus of initial damage: Alzheimer’s, for example, first targets the entorhinal cortex and hippocampus, whereas Parkinson’s disease centers on the substantia nigra (Fu et al. 2018; Kampmann 2024). The same regional specificity is seen in Mendelian disorders as well, which allow us to more readily trace their developmental origins. For example, spinocerebellar ataxia type 1 (SCA1) is caused by expansion of a polyglutamine repeat tract in the gene *ATAXIN-1* (*ATXN1*) (Orr et al. 1993; Banfi et al. 1994). The ATXN1 protein is a transcriptional co-repressor that is expressed throughout the central nervous system (CNS) in both neurons and glia and is also expressed in organs such as the heart and liver (Servadio et al. 1995). In the CNS, it is most abundant in hippocampal pyramidal cells and cerebellar Purkinje cells, yet SCA1 pathology is largely restricted to the Purkinje cells and brainstem (Zoghbi and Orr 1995). Interestingly, a lack of ATXN1 does not cause ataxia but rather learning and memory deficits (Matilla et al. 1998; Crespo-Barreto et al. 2010; Asher et al. 2020): copy number variation of *ATXN1* is associated with Alzheimer’s disease risk (Swaminathan et al. 2011; Swaminathan et al. 2012), and *Atxn1* knock-out in mice potentiates amyloid beta production (Suh et al. 2019). If, however, we delete *Atxn1* and its paralog, *Ataxin-1-*like (*Atxn1l*), the double mutant mice develop quite a different constellation of abnormalities that includes hydrocephalus, omphalocele, lung defects, and perinatal lethality (Lee et al. 2011). Loss of *Atxn1l* by itself causes the same phenotypes but with reduced penetrance (Lee et al. 2011).

It is reasonable to assume that this combination of regional specificity independent of expression level and the very different tissues affected by loss of ATXN1L indicates that ATXN1 and ATXN1L have different native interactors. It was rather surprising, then, to find that they both form a complex with the transcriptional repressor Capicua (CIC) (Lee et al. 2011; Lu et al. 2017), and both proteins are critical for maintaining CIC stability (Lam et al. 2006; Lee et al. 2011; Lu et al. 2017; Wong et al. 2020).

ATXN1 and ATXN1L share a conserved CIC-binding domain (Mizutani et al. 2005; Lam et al. 2006; Kim et al. 2013) and can substitute for each other in binding to CIC, exhibiting partial functional redundancy (Bowman et al. 2007; Crespo-Barreto et al. 2010; Lee et al. 2011). CIC is also broadly expressed (Lee et al. 2011; Kim et al. 2015; Lu et al. 2017) and interacts with ATXN1 throughout the brain (Lu et al. 2017); the effect of the polyglutamine expansion in mutant ATXN1 is to intensify the effect of this interaction, leading to hyper-repression of CIC targets (Fryer et al. 2011; Coffin et al. 2023). The formation of the mutant ATXN1–CIC complex is necessary for Purkinje cell pathology in SCA1 (Rousseaux et al. 2018; Coffin et al. 2023). On the other hand, loss of ATXN1 reduces CIC levels, leading to derepression of CIC targets *Etv4* and *Etv5* (Suh et al. 2019). These transcription factors have binding sites upstream of *Bace1* and increase its expression, which in turn promotes amyloidogenic processing of amyloid precursor protein (APP) and may be a risk factor for Alzheimer disease (Suh et al. 2019). Notably, this upregulation occurs only in the cortex and hippocampus, not in the cerebellum (Suh et al. 2019).

CIC has broad developmental roles in both the peripheral tissues and the CNS. In mice, complete loss of CIC causes perinatal lethality, with hydrocephalus, omphalocele and impaired lung development (Lee et al. 2011; Simon-Carrasco et al. 2017)—reminiscent of *Atxn1l* knockout (KO) mice. In humans, heterozygous loss-of-function variants cause CIC haploinsufficiency syndrome (CHS), a neurodevelopmental disorder characterized by intellectual disability, autism spectrum disorder (ASD), attention deficit hyperactivity disorder (ADHD), and seizures (Lu et al. 2017). Dual loss of both ATXN1 and ATXN1L phenocopies the complete loss of CIC (Lee et al. 2011), and conditional KO in the brain causing a CHS-like phenotype (Lu et al. 2017). How do these two paralogs, both acting through CIC, produce such distinct loss-of-function outcomes?

We hypothesized that the answer might arise from the fact that CIC exists in two isoforms generated by alternative promoter usage, namely, CIC-L (long) and CIC-S (short) (Lam et al. 2006). Both isoforms share the HMG box and C1 DNA-binding domains as well as the ATXN1/ATXN1L-binding domain (ATXN1/1L-BD) (Lam et al. 2006; Jimenez et al. 2012; Lee 2020). The functions of the isoform-specific N-termini remain largely undefined, leaving open the possibility that the two isoforms have distinct biological roles. To determine whether isoform-specific functions contribute to the regional specificity of ATXN1 and ATXN1L gain- or loss-of-function phenotypes, we generated CIC isoform-specific KO mice, characterized their phenotypes, and evaluated transcriptomic changes. We then performed biochemical studies to assess the formation of different complexes. We found that, surprisingly, subtle differences in the relative abundance of CIC isoforms and ATXN1 paralogs at the protein level, together with the differential complexes they assemble, can dictate regional vulnerability.

## RESULTS

### CIC isoforms and ATXN1/ATXN1L show different spatiotemporal expression patterns

To investigate how the expression patterns of CIC isoforms relate to that of the CIC–ATXN1/1L complex, we profiled protein levels of each CIC isoform together with ATXN1 and ATXN1L in the cortex, cerebellum, and lung, of wild-type (WT) mice at four developmental stages: birth (P0), postnatal day 6 (P6), postnatal day 21 (P21), and 12 weeks (**Fig. 1A, B**). These regions and developmental timepoints were selected based on prior evidence of tissue-specific changes due to disruption of the CIC–ATXN1/1L complex: in the forebrain, *Cic* knockout (KO) or *Atxn1/Atxn1l* double KO in mice leads to selective reduction of cortical layers 2–4 beginning around postnatal day 5 (P5), with behavioral abnormalities emerging by 3 months of age (Lu et al. 2017). In the lung, *Atxn1l* KO in mice disrupts the transcriptional program of early alveolarization at P6 and results in reduced survival by P21 (Lee et al. 2011). The cerebellum, which is highly sensitive to ATXN1 gain-of-function toxicity, was included as a comparator to these loss-of-function–vulnerable tissues (Rousseaux et al. 2018).

**Figure 1.**
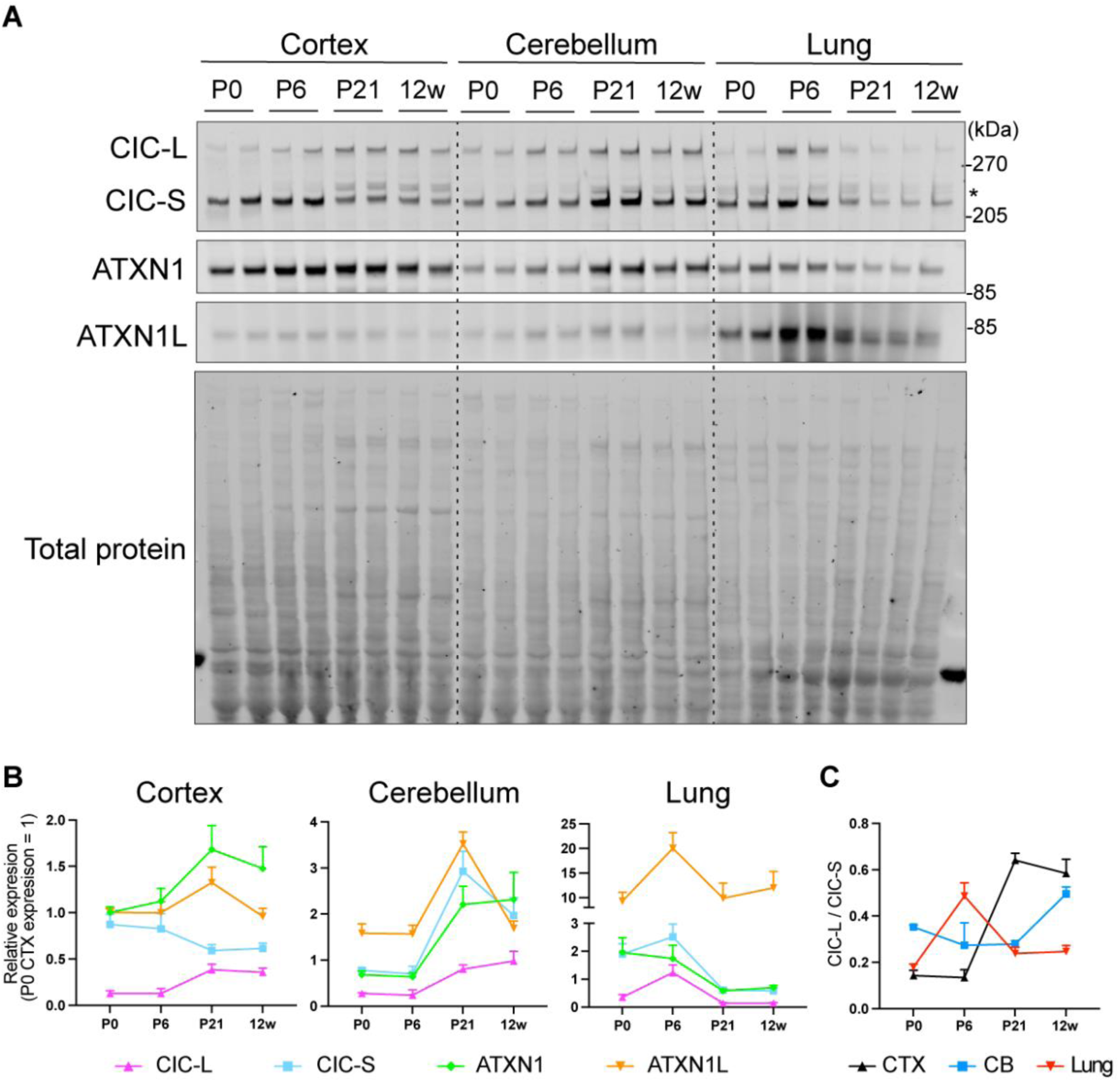
CIC-L and CIC-S show different spatiotemporal expression patterns in mouse cortex, cerebellum, and lung. (A) Representative immunoblot (IB) of CIC-L, CIC-S, ATXN1 and ATXN1L in wild-type (WT) mouse cortex, cerebellum and lung lysates at indicated time points. Antibody against the C terminus of CIC recognizes both isoforms of CIC. P, postnatal day; w: week-old; *, truncated CIC-L product. (B) Quantification of CIC-L, CIC-S, ATXN1, and ATXN1L protein levels. Each protein was normalized to total protein, with each time point normalized to P0 cortex (CTX). CIC isoforms were normalized to total CIC at P0 CTX. Data are mean ± SEM; n = 4 per time point. (C) Ratio of CIC-L to CIC-S at each time point for each region. CTX, cortex; CB, cerebellum.

ATXN1L and both CIC isoforms showed a similar pattern of expression in the lungs, with the highest levels at P6 and decreasing at later timepoints, whereas ATXN1 expression decreases over time after P0 (**Fig. 1A, B**). Notably, ATXN1L levels are considerably higher in the lungs compared to the cortex and cerebellum, with peak expression at P6. In the cortex, CIC-L and CIC-S expression patterns diverged after P6, as reflected in a marked increase in the CIC-L/CIC-S ratio: CIC-L showed a slight increase at P21 that was maintained at 12 weeks, while CIC-S markedly decreased between P6 and P21 and remained low at 12 weeks (**Fig. 1A-C**). In the cerebellum, all four proteins showed a pronounced increase from P6 to P21, followed by a decrease in CIC-S and ATXN1L at 12 weeks. In 12-week-old brains, we also looked at the hippocampus, striatum, and brainstem and found that both CIC isoforms were most highly expressed in the cerebellum and least expressed in the brainstem (**Supplemental Fig. S1**).

Thus, CIC, ATXN1, and ATXN1L show distinct tissue- and isoform-specific expression patterns in mice from birth through young adulthood.

### CIC-S is critical for survival and for lung alveolarization

To dissect the individual roles of each CIC isoform, we used CRISPR/Cas9 to generate *Cic-L*- and *Cic-S*-specific KO mice by targeting the unique first coding exon of each isoform. For *Cic-S-*KO, we targeted *Cic-S* exon 1 with a single guide RNA (sgRNA) and selected founder lines with deletions that removed the in-frame downstream ATG (**Fig. 2A, Supplemental Fig. S2A**, **B**). For *Cic-L*-KO, we used two sgRNAs to introduce a deletion within exon 2, resulting in a frameshift mutation and selective disruption of the long isoform. We confirmed the deletions by Sanger sequencing (**Supplemental Fig. S2C**) and used western blot to validate complete loss of either CIC-L or CIC-S in homozygous KO animals (**Fig. 2B**). Interestingly, CIC-S protein levels are increased in *Cic-L*-KO, whereas loss of CIC-S does not produce a comparable increase in CIC-L. CIC has been reported to bind regulatory regions of its own locus (Weissmann et al. 2018), raising the possibility that there are isoform-specific differences in transcriptional autoregulation of the CIC locus.

**Figure 2.**
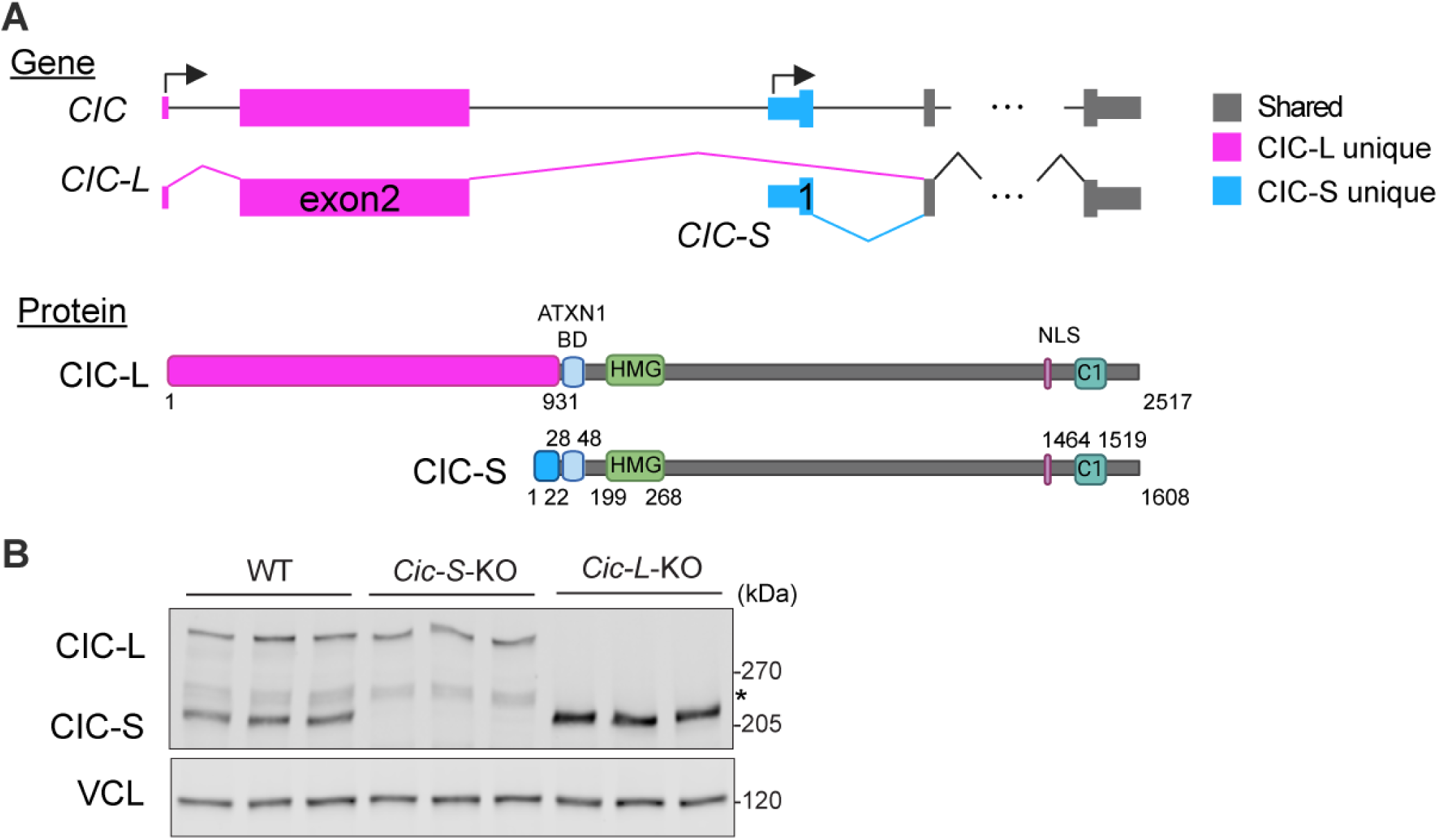
*Cic-L*-KO and *Cic-S*-KO mice were generated using CRISPR. (A) Gene and protein structure of human CIC-L and CIC-S. ATXN1/1L BD, ATXN1/1L binding domain; HMG, HMG box; C1, C1 domain; NLS: Nuclear localization signal (Created in BioRender, https://BioRender.com/kjq2awu). (B) IB of CIC-L and CIC-S of cortex lysate of WT, *Cic-S*-KO and *Cic-L*-KO mice at 1 month of age. Vinculin (VCL) as loading control. n =3 per genotype. *, truncated CIC-L product.

Since germline KO of *Cic* results in perinatal lethality (Lu et al. 2017; Simon-Carrasco et al. 2017), we sought to determine how loss of each isoform affects survival. We used chi-square analysis to compare observed genotype frequencies to the expected Mendelian ratios of WT (25%), heterozygous KO (50%), and KO (25%) at postnatal days 0, 6, and 21 (P0, P6, and P21; **Tables 1** and **2**). Both *Cic-S* and *Cic-L*-KO animals exhibited the expected Mendelian ratios at P0, indicating no embryonic lethality. At P6, neither group showed significant deviation from expected ratios, but by P21, approximately 70% of *Cic-S*-KO had died (chi-square analysis, *p* = 0.007; **Table 1**) while *Cic-L*-KO animals maintained normal survival (chi-square analysis, *p* = 0.74; **Table 2**).

**Table 1.**
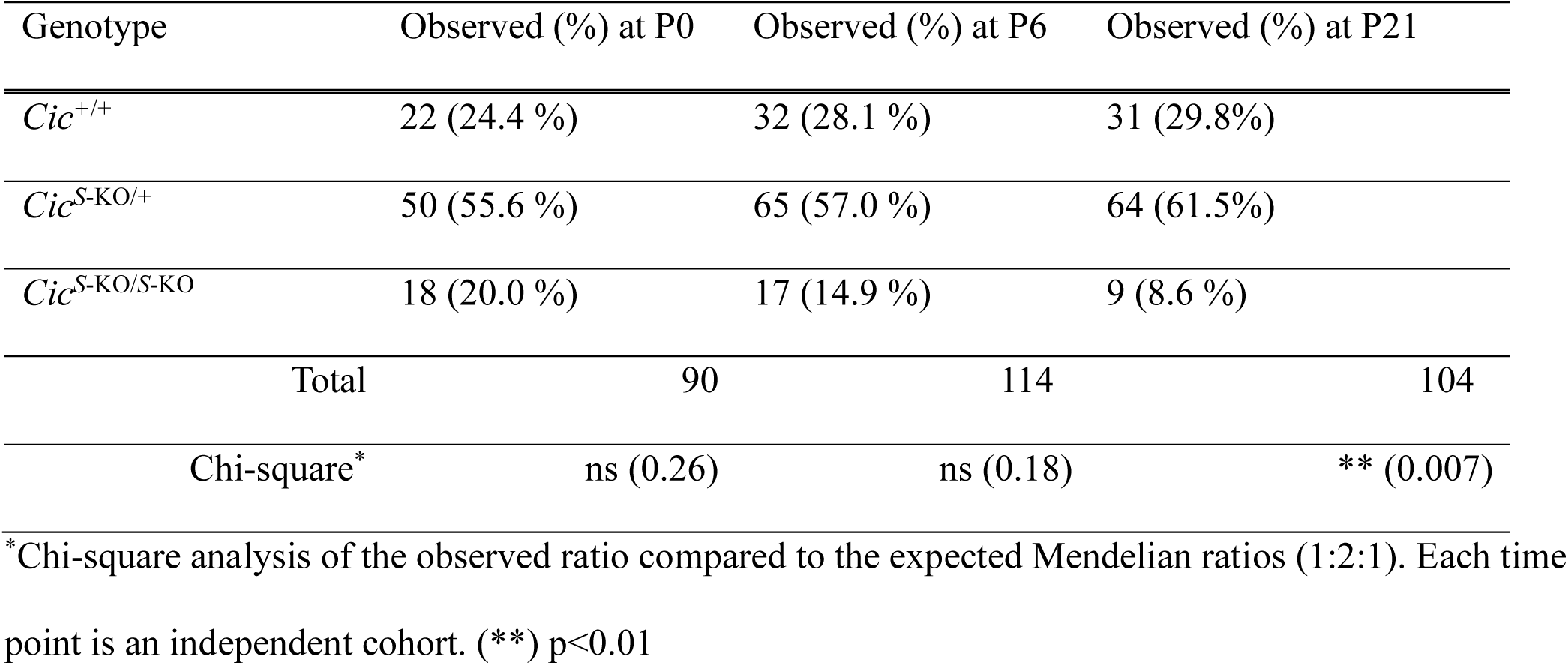
Number of surviving animals from *Cic^S^*^-KO/+^ intercrosses at a given time point.

**Table 2.**
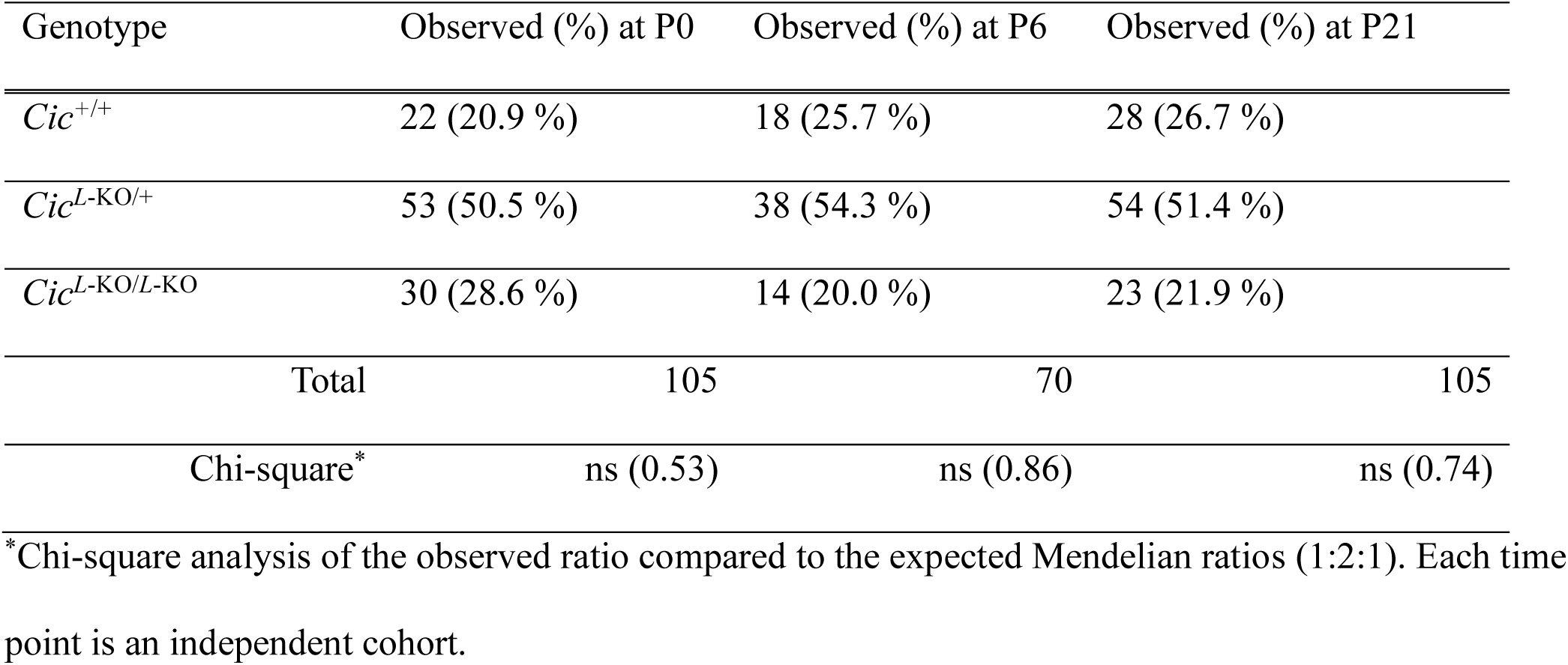
Number of surviving animals from *Cic^L^*^-KO/+^ intercrosses at a given time point.

| Genotype | Observed (%) at P0 | Observed (%) at P6 | Observed (%) at P21 |
| --- | --- | --- | --- |
| *Cic<sup>+/+</sup>* | 22 (20.9 %) | 18 (25.7 %) | 28 (26.7 %) |
| *Cic<sup>L-KO/+</sup>* | 53 (50.5 %) | 38 (54.3 %) | 54 (51.4 %) |
| *Cic<sup>L-KO/L-KO</sup>* | 30 (28.6 %) | 14 (20.0 %) | 23 (21.9 %) |
| Total | 105 | 70 | 105 |
| Chi-square* | ns (0.53) | ns (0.86) | ns (0.74) |

\*Chi-square analysis of the observed ratio compared to the expected Mendelian ratios (1:2:1). Each time point is an independent cohort.

We measured the weight and lifespan of surviving animals beyond P21. *Cic-S*-KO and *Cic-L*-KO mice weighed less than WT throughout the 9-week period, although *Cis-S*-KO stopped gaining weight around week 7 while *Cic-L*-KO mice continued to grow, reaching near WT weight by week 9 (**Fig. 3A**). *Cic-S*-KO mice exhibited significant early lethality after 3 weeks, with 60% (15 of 25 animals) dying before 20 weeks, whereas *Cic-L*-KO animals did not display reduced survival (**Fig. 3B**). Among the *Cic-S*-KOs that died, 80% developed hydrocephalus accompanied by kyphosis, emaciation, and lethargy prior to death. One animal experienced spontaneous and ultimately fatal home-cage seizures (**Fig. 3C**). The remaining two animals that died without clear hydrocephalus or seizures showed further weight loss beyond their already reduced baseline prior to death. Some *Cic-S*-KO mice survived up to 40 weeks without any of these features, however, suggesting incomplete penetrance. Neither *Cic-L*-KO nor WT animals developed any noticeable phenotype during this period.

**Figure 3.**
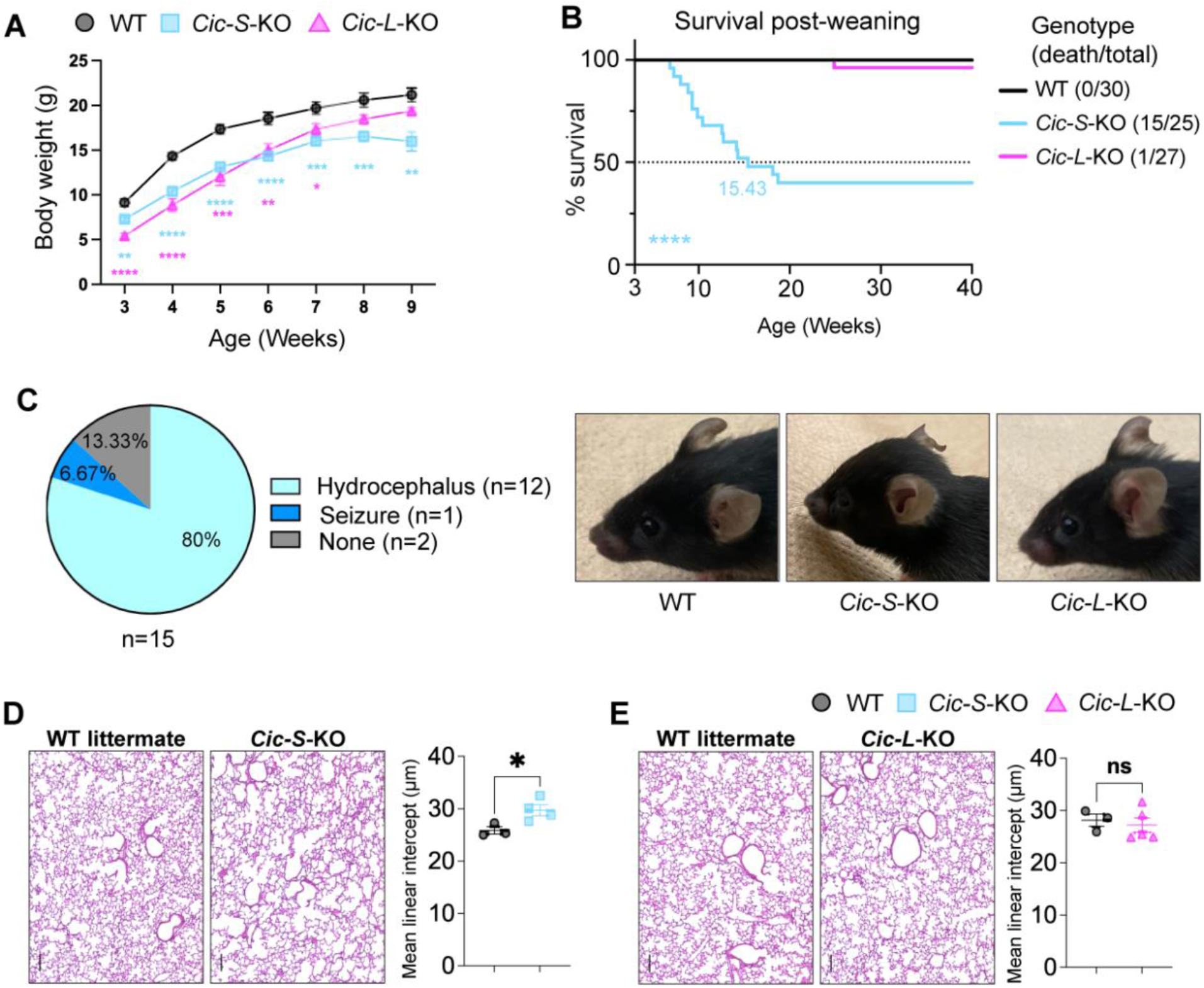
*Cic-S*-KO mice display early lethality and lung alveolarization defects. (A) Weekly body weights of *Cic-S*-KO, *Cic-L*-KO, and their WT littermate from week 3. Data are mean ± SEM; n = 16 per genotype (8 males, 8 females). Two-way repeated-measures ANOVA with Dunnett’s multiple comparisons; (*) p < 0.05, (**) p < 0.01, (***) p < 0.001, (****) p < 0.0001. (B) Kaplan–Meier survival curves post-weaning (3 weeks) for WT, *Cic-S*-KO, and *Cic-L*-KO mice (n = 30, 25, and 27). Mantel–Cox test; (****) p < 0.0001. (C) Left: Phenotypes observed before lethality of *Cic-S*-KO animals (n = 15). Right: Representative photographs of the heads of WT, *Cic-S*-KO and *Cic-L*-KO mice at 9 weeks; *Cic-S*-KO mice show dome-shaped heads due to hydrocephalus. (D,E) Lung tissues from *Cic-S*-KO (D) and *Cic-L*-KO (E) mice and their WT littermates collected at P21. Left: Representative hematoxylin and eosin (H&E)-stained lung sections. Scale bar, 100 μm. Right: Quantification of alveolar airspace by mean linear intercept (MLI). Data are mean ± SEM; n = 3–5 per group. Unpaired t-test; (*) p < 0.05, (ns) p > 0.05.

These abnormalities of the *Cic-S*-KO mice are strongly reminiscent of those previously reported in *Atxn1l*-KO mice (hydrocephalus with incomplete penetrance, lower body weight compared to WT controls, and reduced survival by P21) (Lee et al. 2011). Because *Atxn1l*-KO mice also have impaired alveolarization mediated by loss of CIC function that dysregulates extracellular matrix (ECM) remodeling and alveolarization during late alveolarization stage (P17–P23), which was also present in another line of mice bearing a *Cic* loss-of-function allele (Lee et al. 2011), we performed hematoxylin and eosin (H&E) staining of lung tissue at P21 and quantified the mean linear intercepts (MLI). Despite potential survivor bias in *Cic-S*-KO animals at this stage, the MLI values were significantly greater in *Cic-S*-KO lungs than in WT littermates (*Cic-S*-KO, 29.73 ± 1.05 vs. WT littermate, 25.86 ± 0.71; p=0.038, unpaired t-test; **Fig. 3D**) whereas MLI values of *Cic-L*-KO lungs were comparable to WT littermate controls (*Cic-L*-KO, 27.21 ± 1.36 vs. WT littermate, 28.14 ± 1.17; p=0.658, unpaired t-test; **Fig. 3E**). Therefore, loss of CIC-S, but not CIC-L, impairs alveolarization, supporting functional overlap between CIC-S and ATXN1L.

### CIC-L is crucial for neurobehavioral function

Given the critical role of CIC in neurodevelopment (Lu et al. 2017; Ahmad et al. 2019; Hwang et al. 2020), we performed a battery of behavioral assays in *Cic-S*-KO, *Cic-L*-KO, and WT littermates to assess anxiety-like behavior, general locomotor activity, motor coordination, and social interaction—phenotypes previously linked to CIC dysfunction in both people with CHS and in mouse models (Lu et al. 2017; Tan and Zoghbi 2019). *Cic-L*-KO mice displayed reduced anxiety-like behavior in the elevated plus maze assay, spending significantly more time in the open arms (**Fig. 4A**), and hyperactivity in the open field test, with increased total distance traveled and higher speed compared to controls (**Fig. 4B**). They had a shorter latency to fall from the rotarod (**Fig. 4C**) and took more time to descend in the pole test (**Fig. S3A**). In the fear-conditioning assay, *Cic-L*-KO mice showed deficits in learning and memory, freezing less in both contextual (**Fig. 4D**) and cued fear tests (**Fig. 4E**). In the three-chamber assay, both KO lines preferred to interact with the conspecific than the object (**Fig. S3B**).

**Figure 4.**
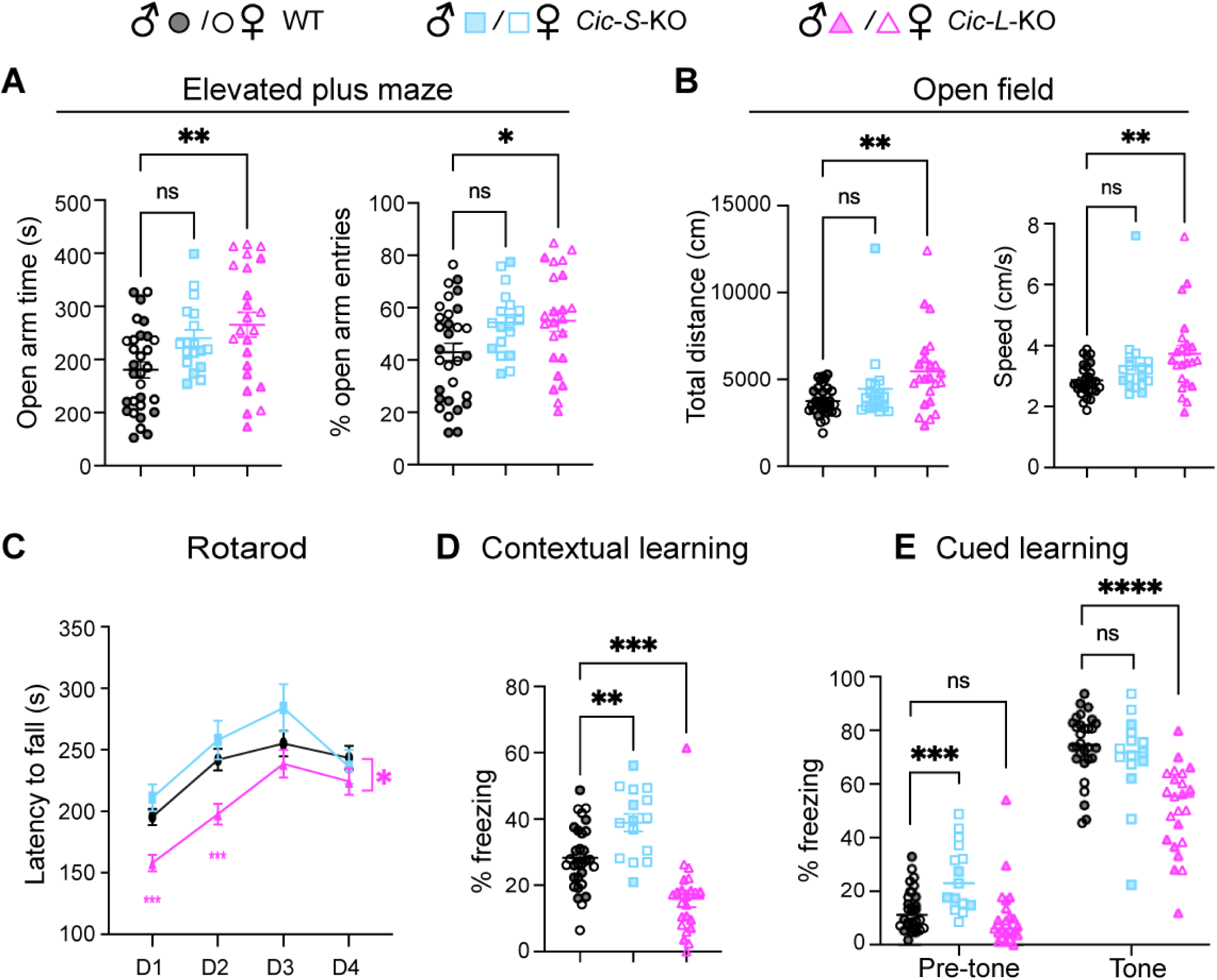
*Cic-L*-KO mice display neurobehavioral deficits. (A-E) Behavior characterization of *Cic-S*-KO, *Cic-L*-KO, and their WT littermate controls. A) Anxiety-like behavior measured using an elevated plus maze. (B) Activity measured with open field assay. (C) Motor coordination measured with rotarod. D, day; males and females combined. (D,E) Learning and memory assessed with fear conditioning assay. For (A,B), n = 30 WT (14 males, 16 females), n = 18 *Cic-S*-KO (5 males, 13 females), and n = 23 *Cic-L*-KO (12 males, 11 females). Due to early lethality of *Cic-S*-KO mice, n = 17 (5 males, 12 females) and n = 15 (4 males, 11 females) *Cic-S*-KO mice were analyzed for (C) and (D,E), respectively. One-way ANOVA with Dunnett’s multiple comparisons for (A,B,D,E); two-way repeated-measures ANOVA with Dunnett’s multiple comparisons for (C). Data are mean ± SEM; (*) p < 0.05, (**) p < 0.01, (***) p < 0.001, (****) p < 0.0001, (ns) p > 0.05.

*Cic-S*-KO mice exhibited minimal behavioral abnormalities (**Fig. 4A-C**). The only significant phenotype that *Cic-S*-KO mice displayed was increased freezing in the contextual test (**Fig. 4D**) and pre-tone in the cued test (**Fig. 4E**), which do not signify any learning or memory deficit. These data suggest that, for the animals that survive because of decreased penetrance, CIC-S is not critical for most neurobehavioral functions, with only mild effects on baseline freezing behavior.

*Cic-L*-KO mice phenocopied *Atxn1*-KO mice, which show learning deficits (Matilla et al. 1998; Crespo-Barreto et al. 2010; Asher et al. 2020). Collectively, the survival and neurobehavioral data point to a functional and spatio-temporal convergence between CIC-L and ATXN1 on the one hand, and CIC-S and ATXN1L on the other.

### Region-specific transcriptomic overlap between CIC isoform and ATXN1 paralog KOs

To determine whether phenotypic similarities between these two pairs of mouse lines reflect shared transcriptional changes, we performed bulk RNA sequencing (RNA-seq) on tissues linked to the phenotype of each CIC isoform loss (P6 lung and 12-week cortex) and compared the differentially expressed genes (DEGs) previously identified in *Atxn1*-KO and *Atxn1l-*KO mice. We used the cerebellum as an internal control, since it does not contribute to the CHS phenotype despite its great sensitivity to polyglutamine-expanded ATXN1 (Rousseaux et al. 2018)

In the P6 lung, *Cic-S*-KO samples exhibited a greater number and magnitude of DEGs (*p*-adjusted < 0.05 and |log2 fold change (Log2FC)| > 0.25) than *Cic-L*-KO, with enrichment of upregulated genes consistent with CIC’s role as a transcriptional repressor (**Supplemental Fig. S4A**, **B**). These differences aligned with more pronounced survival and developmental phenotypes in *Cic-S*-KO animals (**Table 1**, **Fig. 3**). Gene set enrichment analysis (GSEA) (Subramanian et al. 2005) revealed distinct GO term enriched for each isoform KO, with *Cic-S*-KO enriched for immune-related terms (**Supplemental Fig. S4C**) and *Cic-L*-KO for chromosome segregation-related terms (**Supplemental Fig. S4D**).

Consistent with the phenotypic similarities, the *Cic-S*-KO DEGs showed stronger overlap with the *Atxn1l*-KO DEGs (Lee et al. 2011) (Fisher’s exact test, OR = 19.85, *p* = 6.32 × 10⁻¹³²; **Supplemental Fig. S5A**) than *Cic-L*-KO (OR = 3.58, *p* = 1.7 × 10⁻¹²; **Supplemental Fig. S5B**), with most shared genes showing concordant expression changes. Interestingly, although *Atxn1*-KO lungs had only 34 DEGs, just over a third overlapped with the *Cic-S*-KO DEGs (OR = 4.74, p = 5.83 × 10⁻⁵; **Supplemental Fig. S5C**) or the *Cic-L*-KO DEGs (Fisher’s exact test, OR = 9.86, *p* = 8.54 x 10^-8^; **Supplemental Fig. S5D**). Given the role of the *Etv–Mmp* axis in *Atxn1l*-KO alveolarization defects (Lee et al. 2011) and the fact that *Etv1/4/5* are direct CIC targets (Dissanayake et al. 2011; Lee et al. 2011; Weissmann et al. 2018; Coffin et al. 2023), we examined their expression specifically. *Etv4, Mmp8, Mmp9,* and *Mmp12* were strongly upregulated in *Cic-S*-KO lungs, while *Etv4, Mmp12* and *Mmp13* showed mild upregulation in *Cic-L*-KO (**Supplemental Fig. S5E**). GSEA revealed enrichment of extracellular matrix (ECM) assembly-related pathways in *Cic-S*-KO, whereas *Cic-L*-KO showed weak or non-significant enrichment (**Supplemental Fig. S5F**). These results suggest that CIC-S and ATXN1L regulate similar genes in the lung during lung development.

In the 12-week-old cortex, there were more DEGs in the *Cic-L-*KO with larger fold-changes than in *Cic-S*-KO (**Supplemental Fig. S6A, B**). Whereas *Cic-S*-KO top GSEA terms were mostly related to glia or to bioenergetics (**Supplemental Fig. S6C, D**), *Cic-L*-KO terms involved synapse and neurotransmission-related functions (**Supplemental Fig. S6E, F**). Comparison with *Atxn1*-KO cortex DEGs (*unpublished data*) revealed greater overlap and stronger enrichment with *Cic-L*-KO than *Cic-S-*KO, with more genes changing in the same direction (**Supplemental Fig. S7A, B, Supplemental Table S1, 2**). Enrichment analysis of these concordant DEGs showed that only *Cic-L-*KO shared synapse- and memory-related GO terms with *Atxn1*-KO (**Supplemental Fig. S7C, D**). Consistent with previous findings that the *Etv4/Etv5–Bace1* axis is upregulated in *Atxn1*-KO cortex (Suh et al. 2019), we observed similar upregulation of this axis in *Cic-L*-KO cortex (**Supplemental Fig. S7E**). These findings support the idea that CIC-L and ATXN1 regulate similar gene networks that underlie learning and memory.

In the cerebellum, a region generally more resistant to the loss of the CIC–ATXN1/1L complex but clearly vulnerable to the gain of CIC-ATXN1 function (Rousseaux et al. 2018; Coffin et al. 2023), we observed fewer DEGs in *Cic-L*-KO compared to the cortex, but more DEGs in *Cic-S-*KO relative to its cortical counterpart (**Supplemental Fig. S8A, B, Supplemental Table S3, 4**), potentially reflecting higher CIC-S expression in this region at 12 weeks (**Supplemental Fig. S1**). The two KOs also showed distinct GO term enrichments (**Supplemental Fig. S8C–F**), and DEG overlap with *Atxn1*-KO was stronger for *Cic-S*-KO (Fisher’s exact test, OR = 2.41, *p* = 1.69 x 10^-17^; **Supplemental Fig. S9A**) than for *Cic-L*-KO (Fisher’s exact test, OR = 1.61, *p* = 2.85 x 10^-8^; **Supplemental Fig. S9B**). *Etv4*, which was strongly upregulated in the cortex of *Cic-L*-KO, showed less upregulation in the cerebellum, whereas *Etv5* and *Bace1* remained unchanged (**Supplemental Fig. S9C**).

CIC-L and CIC-S, therefore, largely regulate different biological processes. Their respective transcriptional convergences with ATXN1/1L are both region- and timepoint-dependent.

### CIC-S preferentially associates with ATXN1L, and CIC-L associates with ATXN1, in a region-specific manner

Given the region-specific transcriptomic overlap—between the cortex in *Cic-L*–KO and *Atxn1*–KO mice, and the cerebellum in *Cic-S*–KO and *Atxn1*–KO mice—we sought to determine whether CIC isoform–paralog pairs form distinct complexes across brain regions. We performed size exclusion chromatography (SEC) on protein extracts from cortical and cerebellar tissue harvested from 2–3-month-old *Cic-S*-KO and *Cic-L*-KO mice to examine the elution profiles of CIC, ATXN1, and ATXN1L.

In the WT cortex both ATXN1 and ATXN1L co-eluted with CIC in a high molecular weight complex (∼1.7 MDa, fraction 12), as well as in distinct smaller complexes (ATXN1 peaking at fraction 15, ∼300 kDa; ATXN1L at fractions 16–17, ∼170–88 kDa; **Fig. 5A, B**), consistent with prior reports (Lu et al. 2017). In contrast, both *Cic-S*-KO and *Cic-L*-KO cortices showed less ATXN1 and ATXN1L in the high molecular weight fractions and a concomitant increase in smaller fractions, suggesting that CIC is essential for maintaining their incorporation into the larger complex (**Fig. 5A, B**). Interestingly, in *Cic-L*-KO cortex, ATXN1 showed a stronger shift from the large to small complex (large-to-small complex ratio: 0.41± 0.05, vs. WT 1.00± 0.08; *p* = 0.003, one-way ANOVA) than in *Cic-S-*KO (0.67± 0.08; *p* =0.039) (**Fig. 5C**, left), indicating preferential association of ATXN1 with CIC-L. In contrast, in *Cic-S*-KO cortex ATXN1L shifted markedly to smaller fractions (0.11± 0.03, vs. WT 1.00± 0.03; *p* = 0.0001) and was nearly absent from the large complex. This shift was less pronounced in *Cic-L-*KO (0.51± 0.13; *p* = 0.006; **Fig. 5C**, right), suggesting a stronger association of ATXN1L with CIC-S. This redistribution does not appear to be driven by changes in total protein levels: ATXN1 is lower in *Cic-S*-KO compared to WT, yet the reduction in the large complex is greater in *Cic-L*-KO. ATXN1L levels are not significantly different in either isoform knockout compared to WT (**Supplemental Fig. S10A**). These findings confirm isoform-specific associations within the CIC–ATXN1/ATXN1L complex in the cortex.

**Figure 5.**
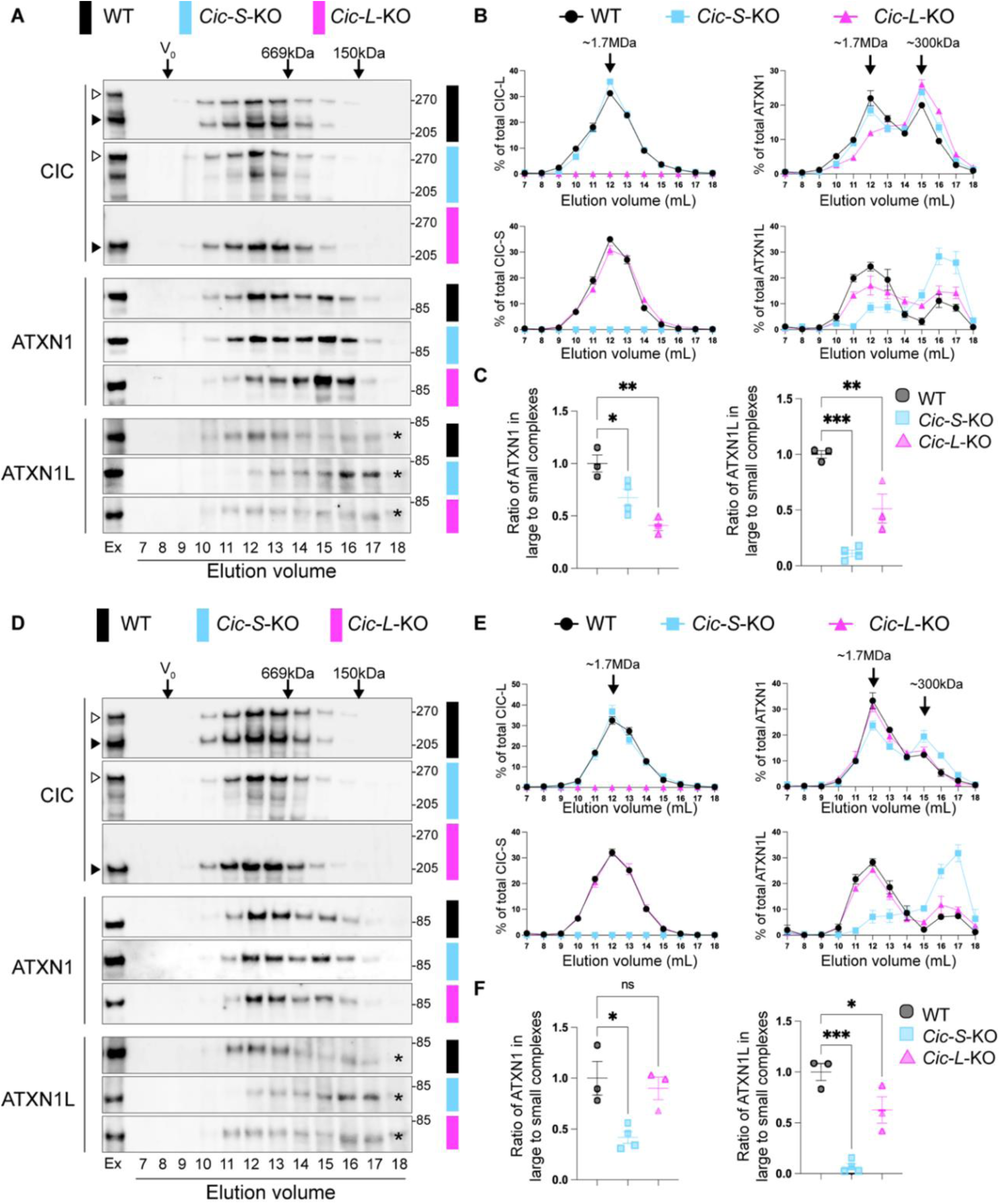
Differential complex formation of ATXN1 and ATXN1L in *Cic-S*-KO and *Cic-L*-KO in the cortex and the cerebellum. (A–C, D–F) Size exclusion chromatography (SEC) analysis of WT, *Cic-S*-KO, and *Cic-L*-KO cortex 877 (A–C) and cerebellum (D–F). (A,D) Representative SEC IBs of cortex (A) and cerebellum (D) lysates probed for CIC-L, CIC-S, ATXN1, and ATXN1L. Void volume (V₀), gel-filtration standards (thyroglobulin, 669 kDa; ADH, 150 kDa), and elution volume (ml) are indicated. Ex, extract; open arrowhead, CIC-L; closed arrowhead, CIC-S; *: a nonspecific band which is observed in *Atxn1l*-KO. (B,E) SEC elution profiles of indicated proteins in cortex (B) and cerebellum (E). The percentage of each protein in each fraction was determined by densitometry and normalized to total signal across fractions for each genotype (n = 3–4 independent extracts). (C,F) Ratio of ATXN1 and ATXN1L in large versus small molecular weight fractions in cortex (C) and cerebellum (F). Left: ATXN1 large (fractions 11–13) to small (fractions 15–17). Right: ATXN1L large (fractions 11–13) to small (fractions 16–17). Ratios were normalized to WT. Data are mean ± SEM. One-way ANOVA with Dunnett’s multiple comparisons; (*) p < 0.05, (**) p < 0.01, (***) p < 0.001.

In the *Cic-S*-KO cerebellum, ATXN1L shifted markedly toward smaller fractions (0.07± 0.03, vs. WT 1.00± 0.08; *p* = 0.0002). The shift was more modest in *Cic-L*-KO (0.63± 0.13; *p* = 0.042; **Fig. 5D–F**), consistent with its preferential association with CIC-S observed in the cortex. In contrast to the cortex, ATXN1 also shifted in *Cic-S*-KO cerebellum (0.42± 0.06, vs. WT 1.00± 0.17; *p* = 0.015), while its distribution remained largely unchanged in *Cic-L*-KO (0.90± 0.11; *p* = 0.82; **Fig. 5D-F**). As in the cortex, changes observed in the cerebellum do not appear to be driven by ATXN1 or ATXN1L levels, as their expression is not significantly altered in either isoform knockout compared to WT **(Supplemental Fig. S10B)**.

The cerebellum has the highest level of CIC-S among all brain regions, as well as the highest total CIC expression (**Supplemental Fig. S1A–D**). ATXN1 is expressed at lower levels, however, resulting in the highest CIC-to-ATXN1 ratio in the cerebellum (**Supplemental Fig. S1E**), which may lead to a greater proportion of ATXN1 bound to CIC. Supporting this hypothesis, comparison of WT cerebellar and cortical SEC patterns (**Supplemental Fig. S11**) revealed a striking difference: the cerebellum exhibits a higher proportion of ATXN1 in the large complex and lower levels in the small complex (**Supplemental Fig. S11C**), a pattern not observed for ATXN1L (**Supplemental Fig. S11D**). The relatively high CIC-S levels may explain why CIC-S is especially important for ATXN1 large complex formation in the cerebellum. Moreover, the combination of a high CIC/ATXN1 ratio likely predisposes the cerebellum to gain-of-function effects of the complex. It is worth noting that cerebellar single-nucleus RNAseq (Kozareva et al. 2021) show that both *Cic* and *Atxn1* are highly expressed in Purkinje cells, which are most vulnerable to SCA1 (**Supplemental Fig. S12**).

Overall, these results indicate that CIC isoforms and ATXN1 paralogs favor particular interactions depending on the context. While the CIC-S–ATXN1L interaction appears strong and consistent across regions, ATXN1 associates with different CIC isoforms in different tissues.

### The unique N-terminal of each CIC isoform mediates the ATXN1 paralog-specific interaction

To understand the mechanism of isoform-specific complex formation of CIC with either ATXN1 or ATXN1L, we first examined the interaction of CIC-L and CIC-S with ATXN1 and ATXN1L by performing coimmunoprecipitation (co-IP) assays in WT cortical lysates. Despite comparable expression levels of CIC isoforms in the cortex, ATXN1 immunoprecipitation showed strong enrichment of CIC-L relative to CIC-S (**Fig. 6A**). Conversely, ATXN1L immunoprecipitation demonstrated an even more pronounced enrichment of CIC-S over CIC-L (**Fig. 6B**), which may explain the more prominent ATXN1L shift observed in SEC (**Fig. 5A-C**). These results support the hypothesis that ATXN1 preferentially interacts with CIC-L, whereas ATXN1L favors CIC-S.

**Figure 6.**
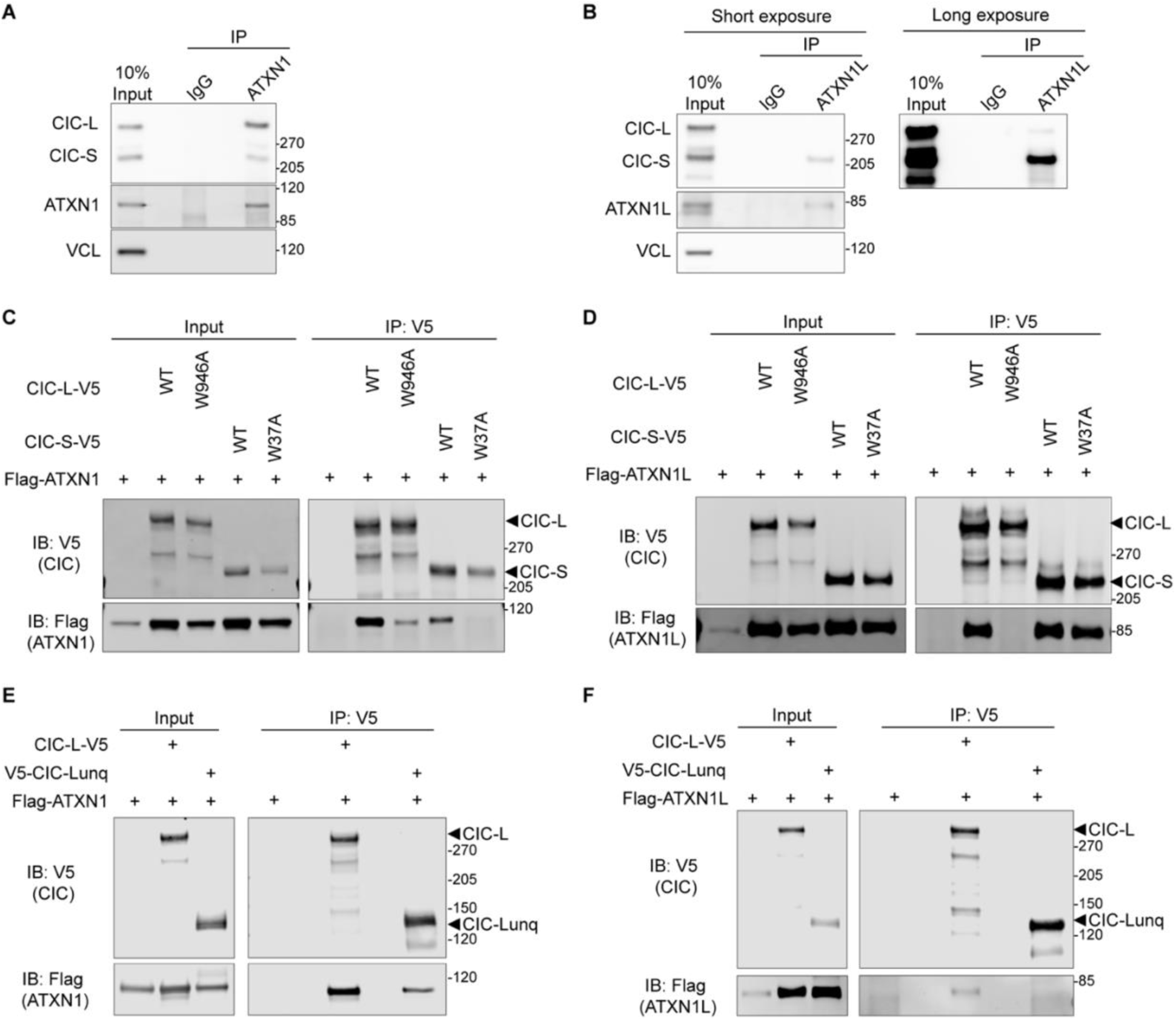
The unique domain of each CIC isoform mediates the preferential interaction with ATXN1 paralogs. (A, B) Representative IBs of immunoprecipitation (IP) experiments from WT cortex. IP of ATXN1 (A) and ATXN1L (B) and co-IPed CIC-L and CIC-S. (C,D) Representative IBs of IP experiments from HEK293T co-transfected with V5-tagged CIC-L or CIC-S, either WT or with an ATXN1/1L binding mutation (W946A or W37A), and Flag-tagged ATXN1 (C) or ATXN1L (D). IP of V5 and co-IPed Flag. (E, F) Representative IBs of IP experiments from HEK293T co-transfected with V5-tagged CIC-L or CIC-L unique N-terminal (CIC-Lunq), and Flag-tagged ATXN1(C) or ATXN1L (D). IP of V5 and co-IPed Flag.

Since CIC-S and CIC-L share a canonical ATXN1/ATXN1L binding domain (ATXN1/1L-BD) that can bind both ATXN1 and ATXN1L, we hypothesized that there should be isoform-specific binding sites outside the ATXN1/1L-BD that mediate preferential interaction between CIC-S and ATXN1L, and CIC-L and ATXN1. To test this possibility, we performed the analysis using human CIC, ATXN1, and ATXN1L sequences. The W37A mutation in CIC-S has been previously shown to disrupt binding with the AXH domain of ATXN1(Kim et al. 2013). Using the corresponding mutation in CIC-L (W946A), we found that CIC-S W37A lost interaction with ATXN1 but CIC-L W946A retained ATXN1 binding (**Fig. 6C**). This suggests that CIC-L contains an additional ATXN1-binding region. We saw the opposite with ATXN1L, despite CIC-S containing only 22 isoform-specific amino acids at its N-terminus: ATXN1L still bound CIC-S W37A but failed to interact with CIC-L W946A (**Fig. 6D**). These data indicate that distinct, noncanonical regions in each isoform mediate selective interactions with ATXN1 and ATXN1L.

To test whether the N-terminal region of CIC-L can mediate the selective interaction with ATXN1, we expressed this region alone and performed co-IP. The CIC-L N-terminus, which spans 931 amino acids and does not contain the ATXN1/1L-BD, interacted with ATXN1 but not with ATXN1L (**Fig. 6E, F**). Thus, the N-terminal region of CIC-L is sufficient to mediate interaction with ATXN1, but not ATXN1L, and may thereby contribute to isoform-paralog specificity.

Together, these findings reveal a previously unrecognized mechanism by which CIC isoforms interact with different ATXN1 paralogs through isoform-specific domains, establishing a novel molecular basis for paralog-specific complex formation.

### CIC-L expression is more dependent on ATXN1 than CIC-S

Given the preferential interaction between ATXN1 and CIC-L, and the known role of ATXN1 in maintaining CIC protein stability (Lam et al. 2006), we hypothesized that CIC-L would be more dependent on ATXN1. To test this, we measured CIC-L and CIC-S protein levels in WT and *Atxn1*-KO mice across brain regions sensitive to ATXN1 loss (cortex, hippocampus) or gain (cerebellum, brainstem). CIC-L levels were significantly lower in all four regions in *Atxn1*-KO mice, but CIC-S levels were more modestly reduced, and only in the cerebellum (**Supplemental Fig. S13A-D**). The cerebellar reduction in CIC-S was consistent with our SEC analysis showing a shift of CIC-S in ATXN1-containing fractions (**Fig. 5D-F**). IP of ATXN1 confirmed a preferential association with CIC-L in all brain regions examined (**Supplemental Fig. S13E**). These findings indicate that the cortical and hippocampal abnormalities in *Atxn1*-KO mice may result primarily from the loss of CIC-L function.

## DISCUSSION

The present study uncovers functionally distinct roles for the two CIC isoforms, CIC-L and CIC- S. The overlap of transcriptomic changes in *Cic-S*-KO and *Atxn1l*-KO mice explains why these two lines show the same early lethality, hydrocephalus, and lung alveolarization defects, while *Cic-L*-KO and *Atxn1-*KO mice share learning and memory deficits. These phenotypic parallels reveal an unexpected degree of specificity in CIC–ATXN1 complex formation, given that both isoforms share the canonical ATXN1/ATXN1L binding domain (ATXN1/1L-BD); the specificity is attributable to the unique N-terminal domains of each isoform that favor binding between CIC-S and ATXN1L, and between CIC-L and ATXN1. The sharp distinctions between the two sets of phenotypes provide compelling *in vivo* support of the idea that different isoforms greatly expand protein interaction capacities and may indeed serve as “functional alloforms” (Yang et al. 2016).

Prior studies in *Drosophila* have shown that CIC isoforms play distinct roles during development: Cic-S is critical for early embryogenesis through its interaction with Groucho (Fores et al. 2015), while Cic-L is essential for oogenesis (Rodriguez-Munoz et al. 2022). These findings support the idea that the divergent N-terminal regions of CIC isoforms confer isoform- and context-specific functions. However, *Drosophila* Cic-S has evolved on its own (Fores et al. 2015), with the species possessing only Atx-1 (containing the AXH domain) and lacking the paralog pairs found in mammals. Furthermore, there is no evidence for an endogenous functional complex between *Drosophila* Cic and Atx-1. By using a mammalian system in which both CIC isoforms and ATXN1 paralogs are highly conserved, we identified a previously unrecognized layer of regulation: isoform–paralog–specific interactions.

These isoform-specific roles are not only of evolutionary interest but also carry direct clinical relevance. Recently, a CIC-L–specific mutation was reported in a patient with CHS (Sharma et al. 2022), which is consistent with our finding that CIC-L is critical for neurological function. There may be additional patients with CIC-L variants who remain unrecognized. Furthermore, given the role of CIC-L in learning and memory and its regulation of the *Etv4/5–Bace1* axis, CIC-L variants could, like *ATXN1* (Bertram et al. 2008; Bettens et al. 2010; Swaminathan et al. 2011; Swaminathan et al. 2012; Suh et al. 2019). contribute to susceptibility to Alzheimer’s disease. More broadly, the expression of different isoforms can be dysregulated in disease, but gene-level expression measures commonly used in biology average the translational potential of each isoform and therefore may not accurately reflect protein concentration. It has also been proposed that isoform-specific therapeutics would improve efficacy and reduce adverse effects (Kjer-Hansen et al. 2024).

Our findings help resolve two longstanding questions: (1) What underlies the regional vulnerability to loss or gain of the CIC–ATXN1 complex despite its broad expression? (2) How does the loss of ATXN1 versus ATXN1L lead to distinct phenotypes, despite both acting through CIC? We propose that the combination of isoform–paralog–specific interactions and regional expression patterns determines the differential sensitivity to complex disruption. Consistent with this model, the similarity in learning and memory deficits, along with upregulation of the *Etv4/5–Bace1* axis, in both *Atxn1*-KO and *Cic-L*-KO mice suggests that these phenotypes arise from preferential CIC-L–ATXN1 complex formation in the cortex and possibly the hippocampus.

Although we did not assess isoform-specific contributions to the cerebellar and brainstem pathology seen in SCA1, the current study does shed light on this question. We find that total CIC levels and CIC/ATXN1 ratios are highest in the cerebellum, with a greater proportion of ATXN1 being incorporated into the large complex. Our prior work in SCA1 mice demonstrated that ∼50% reduction of total CIC (affecting both isoforms) (Fryer et al. 2011) or a polyQ-ATXN1 mutation that would abolish binding to both isoforms fully rescues the Purkinje cell phenotype (Rousseaux et al. 2018; Coffin et al. 2023). These findings indicate that total CIC levels bound to ATXN1, rather than isoform identity, are critical for Purkinje cell degeneration. By contrast, CIC expression is lowest in the brainstem, which may be why loss of the ATXN1-CIC interaction does not fully rescue brainstem SCA1 phenotypes (Fryer et al. 2011; Coffin et al. 2023) and why CIC motifs are less enriched in the brainstem’s disease- associated gene network (Friedrich et al. 2018). In SCA1, the brainstem’s ATXN1 protein complex composition is distinct from that in the cerebellum (Jafar-Nejad et al. 2011), indicating that brainstem pathology arises through different mechanisms.

Our findings also allow us to draw some inferences that can be tested in the future. First, ATXN1 loss does not fully recapitulate CIC-L deficiency: transcriptomic comparisons reveal only partial overlap between *Atxn1*-KO and *Cic-L*-KO datasets, indicating that ATXN1 and CIC-L have functions outside of their shared complex. Indeed, ATXN1 has interactors beyond CIC (Chen et al. 2003; Tsuda et al. 2005; Serra et al. 2006; Lim et al. 2008; Coffin et al. 2023), and such partners may act in compensatory transcriptional or chromatin regulatory pathways, buffering CIC-dependent gene expression programs.

What are the ATXN1/ATXN1L-independent mechanisms that distinguish CIC-L from CIC-S? If isoform differences were driven solely by interactions with ATXN1 and ATXN1L, *Cic-L*-KO and *Cic-S*-KO transcriptomes would be expected to fully overlap, differing only in the magnitude of differential expression. Instead, the partial overlap of DEGs may reflect isoform-specific interactors that influence target gene selection. Alternatively, the isoforms may differ in cell type expression, affecting distinct cellular populations. Divergent GO term enrichments support this notion. Profiling isoform-specific expression across cell types and applying single-cell transcriptomics to each isoform KO will be critical to dissect these contributions.

Finally, our findings highlight a broader principle that paralogous proteins and their isoform-specific partners can form distinct regulatory complexes with completely different biological functions. The CIC–ATXN1/1L complex may serve as a paradigm for understanding how paralog-isoform specific interactions shape regional vulnerability and disease susceptibility in other diseases such as those involving amyloid precursor protein (Kang et al. 1987; Kitaguchi et al. 1988; Ponte et al. 1988; Tanzi et al. 1988; Wasco et al. 1992; Sprecher et al. 1993; Wasco et al. 1993), tau (Wang and Mandelkow 2016; Chung et al. 2021), or α-synuclein (Rhinn et al. 2012), all of which exist within paralog families and have multiple isoforms.

## MATERIALS AND METHODS

### Mouse husbandry

All mice were backcrossed to C57BL/6J background. All mice were housed and maintained in the animal facilities at Baylor College of Medicine (BCM) on a 12-hour light-dark cycle. BCM Institutional Animal Care & Use Committee (IACUC) approved all animal care and experimental procedures.

### HEK293T cell line

Human embryonic kidney cell T (HEK293T) (ATCC, CRL-3216) were cultured in Dulbecco’s modified Eagle’s medium (DMEM, Corning, MT10013CV) supplemented with 10% fetal bovine serum (Gibco A4736201) and 1X antibiotic-antimycotic (Gibco, 15240096). Cell cultures were maintained at 37 °C in a humidified 5% CO_2_ incubator.

### Generation of *Cic-L*-KO and *Cic-S*-KO mice

*Cic-L*-KO and *Cic-S*-KO alleles were generated using CRISPR/Cas9 genome editing. For *Cic-L*-KO, two crRNAs targeting exon 2 of *Cic-L*, the first coding exon unique to the CIC-L isoform, were designed (5′-CGCACCTGGCGCACGACTGC-3′ and 5′-TCCAGCCGCACCGACTGCTG-3′; IDT) to generate a large deletion. For Cic-S-KO, a single crRNA targeting exon 1 of Cic-S was designed (5′-ACGCGGGTATCAGGGGCCTG-3′; IDT).

crRNA, tracrRNA, and Cas9 protein were prepared as described previously (Coffin et al. 2023) with modifications to total reaction volume, concentration and delivery method. Final concentrations were

6.6 μM crRNA, 6.6 μM tracrRNA, and 6 μM Cas9 in T_10_E_0.1_ buffer. For *Cic-L*-KO, each crRNA was diluted to 3.3 μM. The mixture was electroporated into zygotes by the Genetically Engineered Rodent Model (GERM) Core at Baylor College of Medicine (BCM).

Founders were screened by PCR using primers flanking the expected deletion sites (*Cic-L*-KO: 5′-TCCTCCTCTACTGACACGGC-3′, 5′-TAGGACAAGCAGCTCATCACC-3′; *Cic-S*-KO: 5′-GGCCAAGGAGCGGGTTAC-3′, 5′-TCCTAGTGCAGTACCGTCCA-3′). PCR products were TOPO cloned (Invitrogen) and analyzed by Sanger sequencing for frameshift deletions. *Cic-S*-KO founders were selected for deletions eliminating the downstream ATG in exon 1. Founder lines were backcrossed to C57BL/6J WT mice (Jackson Laboratory), and germline transmission was confirmed by sequencing of F1 progeny.

Immunoblotting was performed on homozygous KO animals to confirm the loss of the corresponding CIC isoform. One independent founder line was established for each isoform KO and maintained by successive backcrossing (>5 generations) to C57BL/6J prior to experimental use.

### Genotyping of *Cic-L*-KO and *Cic-S*-KO mice

Genotyping was performed by PCR using allele-specific primers. For *Cic-L*, WT and KO alleles were distinguished using two primer sets: *Cic-L*-WT (5′-ACTGGAGAAGGGAGCTGCAC-3′ and 5′-CTGTCCCTCGAAGTCTCATCG-3′) and *Cic-L*-KO (5′-TCCTCCTCTACTGACACGGC-3′ and 5′- TAGGACAAGCAGCTCATCACC-3′). For *Cic-S*, a single primer pair (5′-GGCCAAGGAGCGGGTTAC-3′ and 5′-TCCTAGTGCAGTACCGTCCA-3′) was used, and WT versus KO alleles were distinguished based on product size.

### Lung Histopathology

The lungs were harvested for morphometry studies on postnatal day (P) 21, as described before (Shrestha et al. 2019). Both lungs of the experimental mouse (n=3-5/group) were inflated at 25 cm pressure for at least 10 minutes via tracheal cannulation with 4% paraformaldehyde. The inflated lungs were then fixed overnight with 4% paraformaldehyde at room temperature with gentle agitation. The fixed lungs were stored in 70% ethanol in a refrigerator until further processing for lung histopathology. The lungs were then paraffin-embedded, sliced into 5 µm thin sections, and stained with H&E. We examined at least five random non-overlapping fields of 10X H&E stained images from both lungs of each animal to quantify alveolarization by mean linear intercepts (MLIs). To calculate the MLI, grids of horizontal and vertical lines were placed on the H&E stained lung images, and the number of times the lines intersected with the lung tissue was counted. The MLI was determined by dividing the total length of the grid lines by the number of intersects.

### Behavioral Assays

All behavioral analyses were conducted during the light phase of a 12-hour light-dark cycle by an experimenter blinded to mouse genotype. Mice had *ad libitum* access to food and water except during testing. Age-matched mice were used in all experiments, and wild-type (WT) littermates from both C*ic-S*-KO and *Cic-L*-KO breeding were pooled as controls. Mice were habituated in the test room for at least 30 min prior to each assay. Tests were performed under room lighting of 150 lux with background white noise at 60 dB.

#### Elevated plus maze

Mice (8-10 weeks of age) were placed in the center of an elevated plus maze consisting of two open arms (25 × 8 cm, with 0.5 cm raised edges) and two closed arms (25 × 8 cm, enclosed by 15 cm high walls), elevated 50 cm above the floor. Behavior was recorded for 10 min using a camera and ANY-maze video tracking system (Stoelting, Wood Dale, IL, USA). Measures included time spent, distance travelled and number of entries in open and closed arms.

#### Open field assay

Mice (9-11 weeks of age) were placed in the center of a 40 × 40 × 30 cm chamber equipped with photobeams (AccuScan, Columbus, OH, USA) and allowed to explore for 30 min testing period. Locomotor activity was automatically analyzed using AccuScan Fusion software (Omnitech). Parameters included total distance traveled and mean motor speed.

#### Rotarod

This assay was performed as described previously (Coffin et al. 2023). Mice (9-11 weeks of age) were placed on a rotating rod (3 cm diameter, 30 cm length; Ugo Basile) and tested for four consecutive days, with four trials per day and a minimum 30-min inter-trial interval. Each trial lasted a maximum of 10 min. The rod accelerated linearly from 5 to 40 rpm over the first 5 min, and then remained at 40 rpm for the final 5 min. Latency to fall was recorded, with two consecutive passive rotations scored as a fall.

#### Conditioned fear

On day 1, mice (19-21 weeks of age) were placed in a conditioning chamber (Actimetrics, Coulbourn instruments, Wilmette, IL) and presented with two conditioned stimulus (CS)-unconditioned stimulus (US) pairings, programmed using the FreezeFrame software (Actimetrics). The trial comprised of 2 min of no sound, followed by 30 s of sound at 80 dB (CS), which was followed by a mild 0.8 mA foot shock for 2 s (US). The tone-shock pairing was repeated after 2 minutes. Twenty-four hours later, contextual memory was assessed by returning the mice to the same chamber for 5 min and quantifying freezing behavior. One hour later, cued fear memory was assessed in a novel chamber by monitoring freezing for 3 min without stimulus, followed by 3 min with the tone. Freezing was recorded and quantified using FreezeFrame software (Actimetrics).

### Protein extraction for size exclusion chromatography

Cortical or cerebellar tissues were collected from two 2–3-month-old mice per genotype and pooled per biological replicate. Tissues were homogenized in TX buffer (50 mM Tris, pH 8.0; 75 mM NaCl; 0.5% Triton X-100; Sigma) supplemented with freshly added protease inhibitor (GenDEPOT, P3100-020) and phosphatase inhibitors (GenDEPOT, P3200-020) using a Dounce homogenizer. Homogenates were extracted at 4 °C for 20 min with rotation and clarified by centrifugation at 20,000 × g for 10 min. Supernatants were re-centrifuged, filtered through 0.45 μm centrifugal filters (Millipore), and protein concentrations were determined by BCA assay (Pierce).

### Size exclusion chromatography

Gel filtration chromatography was performed on a Superose 6 10/300 GL column (Cytiva) using an AKTA Pure system at 4 °C. The column was equilibrated in 50 mM Tris (pH 8.0), 50 mM NaCl, 0.1% Triton X-100. Column calibration was performed with thyroglobulin (667 kDa), apoferritin (440 kDa), alcohol dehydrogenase (150 kDa), BSA (66 kDa), and cytochrome C (14.3 kDa). Fractions (1 mL) were collected, supplemented with protease inhibitor (GenDEPOT, P3100-020) and phosphatase inhibitors (GenDEPOT, P3200-020), and analyzed by SDS-PAGE on NuPAGE 3–8% Tris-Acetate gels (Invitrogen).

### Cloning

The hCIC-S cDNA was obtained from a previously described plasmid (Lam et al. 2006). The unique N-terminal region of hCIC-L was synthesized (GenScript) and cloned into a PU57 backbone. Because direct amplification of the ∼4.8 kb CIC-Shared region was hindered by high GC content at the 3′ end, a two-step cloning strategy was employed: the 5′ and 3′ CIC fragments were first assembled into pENTR223 containing a synthetic multiple cloning site with an AvrII site, followed by insertion of the ∼4 kb central CIC fragment by PCR to generate full-length CIC-Shared.

A W37A point mutation (W37A in CIC-S, corresponding to W946A in CIC-L) was introduced into hCIC-Shared in pENTR223 using gene blocks and Gibson assembly (New England Biolabs (NEB), E2621L) after XhoI (5′ end) and StuI (N-terminal) digestion. CIC-L and CIC-S entry clones were generated by Gibson assembly of the respective unique 5′ regions into the CIC-Shared backbone, followed by addition of C-terminal V5 tags. The hCIC-L–unique sequence was cloned into pENTR223 with an N-terminal V5 tag and a stop codon after amino acid 931. All entry clones were recombined into pcDNA3.1-DEST (derived from pcDNA3.1-3×FLAG-DEST, with stop codons preventing FLAG expression) using Gateway LR recombination (Thermo Fisher).

The hATXN1L cDNA (Horizon Discovery) was cloned with an added N-terminal FLAG tag into pLV-EF1α-IRES-Puro (Hayer et al. 2016) using PCR amplification and Gibson assembly. The FLAG-hATXN1 coding sequence was obtained from the previously described pcDNA1-ATXN1 plasmid (Kim et al. 2013) and cloned into pcDNA3.1 by Gibson assembly.

### Transfection and co-immunoprecipitation in HEK293T cells

HEK293T cells were transfected with the indicated plasmids using TransIT-293 transfection reagent (Mirus Bio) according to the manufacturer’s instructions and incubated for 48–72 h. Cells were washed with PBS and either processed immediately or stored at –80 °C. Frozen cells were thawed on ice and lysed in cold lysis buffer (75 mM NaCl, 5 mM MgCl₂, 50 mM Tris-HCl (pH 8.0), 0.5% Triton X-100, with protease and phosphatase inhibitors) for 30–60 min at 4 °C with rotation. Lysates were cleared by centrifugation at 20,000 × g for 20 min at 4 °C.

For co-immunoprecipitation, lysates from one well of a 6-well plate were incubated with 1 µg V5 antibody (Invitrogen) pre-bound to 15 µL Protein G Dynabeads (Fisher Scientific, Cat. #10-009-D) for 2–4 h at 4 °C with rotation. Beads were washed four times with lysis buffer, and bound proteins were eluted in 1× NuPAGE Sample Reducing Agent and NuPAGE LDS Sample Buffer (Invitrogen), boiled for 10 min, and analyzed by SDS-PAGE on NuPAGE 3–8% Tris-Acetate gels (Invitrogen).

### Immunoprecipitation in mouse brain tissue

Cortices from 12-week-old WT mice were homogenized in either cold NEMT buffer (50 mM Tris-HCl pH 7.5, 150 mM NaCl, 1 mM EDTA, 0.5% NP-40) or lysis buffer (75 mM NaCl, 5 mM MgCl₂, 50 mM Tris-HCl pH 8.0, 0.5% Triton X-100), each supplemented with freshly added protease inhibitor (GenDEPOT, P3100-020) and phosphatase inhibitors (GenDEPOT, P3200-020). Tissue homogenization was performed using an electric pestle (handheld polytron, WPR, 47747-370), followed by mechanical shearing through a 27G syringe. Lysates were incubated on ice for 20 min and clarified by centrifugation at 20,000 × g for 20 min at 4 °C. Ten percent of the supernatant was reserved as input, and the remaining lysate was precleared with Dynabeads Protein G (Fisher Scientific, 10-009-D) for 30 min at 4 °C. Precleared lysates were incubated for 2–3 h at 4 °C with 1–2 µg of antibody—anti-ATXN1 (534, in-house (Gennarino et al. 2018)), anti-ATXN1L (881, in-house), or mouse IgG control—conjugated to 30 µL Dynabeads Protein G. Beads were washed four times with the corresponding lysis buffer and final wash with PBS. Bound proteins were eluted in 1× NuPAGE Sample Reducing Agent and NuPAGE LDS Sample Buffer (Invitrogen), boiled for 10 min, and analyzed by SDS-PAGE on NuPAGE 3–8% Tris-Acetate gels (Invitrogen).

For *Atxn1*-KO experiments, cortex, hippocampus, cerebellum, and brainstem were dissected from 4-week-old WT and *Atxn1*-KO mice and processed using the ATXN1 immunoprecipitation protocol described above.

### Protein extraction for immunoblotting

Brain and lung tissues were collected at postnatal day zero (P0), P6, P21, and 12 weeks and snap-frozen in liquid nitrogen and kept in -80°C until processing. For protein extraction, frozen tissues were homogenized in cold lysis buffer (75 mM NaCl, 5 mM MgCl₂, 50 mM Tris pH 8.0, 0.5% Triton X-100, supplemented with freshly added protease inhibitor (GenDEPOT, P3100-020) and phosphatase inhibitors (GenDEPOT, P3200-020) using an electric pestle (handheld polytron, WPR, 47747-370), followed by mechanical shearing through a 27G syringe. Lysates were incubated on ice for 20 min and centrifuged at 20,000 × g for 20 min at 4 °C. Protein concentrations in the supernatant were measured using the Pierce BCA Protein Assay Kit (Thermo Fisher Scientific), and samples were normalized to equal concentrations.

### Western blot analysis

For immunoblotting, samples were mixed with NuPAGE Sample Reducing Agent and NuPAGE LDS Sample Buffer (Invitrogen), boiled for 10 min, and run on NuPAGE 3–8% Tris-Acetate gels (Invitrogen). Proteins were transferred to nitrocellulose membranes (Amersham, #10600004) using transfer buffer containing 10% methanol. Membranes were blocked with either 5% skim milk in Tris-buffered saline with 0.1% Tween® 20 (TBST) or Intercept (TBS) Blocking Buffer (LI-COR Biosciences) for 1 h at room temperature and incubated overnight at 4 °C with primary antibodies diluted in 3% BSA in TBST. The following primary antibodies were used: anti-CIC (rabbit, in-house, 1:1,000), anti-ATXN1(Servadio et al. 1995) (rabbit, in-house, 11750VII, 1:2,000), anti-ATXN1L (mouse, 674, in-house, 1:1,000), anti-Vinculin (mouse, Sigma-Aldrich, 1:5,000), anti-Flag (Sigma-Aldrich, 1:4000), and anti-V5 (Invitrogen, 1:2,000).

After washing in TBST, membranes were incubated for 1 h at room temperature with either HRP-conjugated secondary antibodies (Jackson ImmunoResearch, 1:10,000 in 5% milk/TBST) or fluorophore-conjugated secondary antibodies (LI-COR Biosciences, 1:10,000 in Intercept Blocking Buffer). HRP blots were developed using ECL (Cytiva, RPN2236) or SuperSignal Femto (Thermo Fisher Scientific, 34095) and imaged with a GE Amersham Imager 680 or ChemiDoc MP Imaging System or LI-COR Odyssey M system. Fluorescent blots were imaged directly using the LI-COR Odyssey M system.

### Statistical analysis

All statistical analyses were performed using GraphPad Prism 9.0, except for RNA-seq analyses. Survival of animals was compared to the expected Mendelian ratio using a chi-square test. Comparisons between two groups were performed using unpaired t-tests. Comparisons involving more than two groups were analyzed by one-way or two-way ANOVA with appropriate post hoc multiple-comparisons tests. Repeated-measures data were analyzed by two-way repeated-measures ANOVA. Survival curves were analyzed using the Mantel–Cox test. Specific statistical tests are indicated in the figure legends. Data are presented as mean ± SEM. In each case, *, **, ***, **** and ns denote p<0.05, p<0.01, p<0.001, p<0.0001 and p>0.05, respectively.

RNA-seq analyses were performed in R using DESeq2, as described in the supplementary methods. Fisher’s exact test was used to calculate odds ratios and p-values for gene set overlaps.

Sample sizes were based on prior studies and standard practice in the field. Biological replicates refer to independent animals. Experimental analyses were performed in a blinded manner whenever possible.

### Data and material availability

All data needed to evaluate the conclusions in this study are present in here and/or the in the Supplemental Material. RNA sequencing reads and processed data were deposited to GEO (GSE310453).

## ACKNOWLEDGEMENTS

We thank Ji-Yoen Kim, Sameer Bajikar, Dah-eun Chloe Chung, Harini Pallikarana, Sih-Rong Wu, and Xue Deng for their critical comments on the manuscript, and Harry Orr and Vicky Brandt for editorial input. We thank Zhijian Yu for technical support in bioinformatics analyses during manuscript revisions.

We further thank the Baylor College of Medicine (BCM) Genetically Engineered Rodent Model (GERM) Core, Jan and Dan Duncan Neurological Research Institute (NRI) Rodent Neurobehavior Core, and Neuropathology Core. Funding: This work was funded by National Institute of Aging F31 AG077918 (H.L.), Cockrell Family Funding (H.L.), National Institute of Neurological Disorders and Stroke R01 NS027699 (H.Y.Z.), Freedom Together Foundation MR-2023-4260 (H.Y.Z.), Howard Hughes Medical Institute (H.Y.Z.), National Heart, Lung, and Blood Institute R01 HL181470 (B.S.), Chao Endowment (Z.L.), Huffington Foundation (Z.L.), National Institutes of Health Cancer Center P30 CA125123 (BCM GERM Core), Cancer Biology Research 1R50CA283804 (BCM GERM Core), Eunice Kennedy Shriver National Institute of Child Health and Human Development Intellectual and Developmental Disabilities Research Center P50HD103555 (NRI Rodent Neurobehavior Core and Neovascularization Core).

## Author contributions

H.L. and H.Y.Z. conceived the study, designed the experiments, and analyzed and interpreted the data. H.L. generated mouse models. H.L., E.V.G., and E.M.R. performed protein biochemistry from mouse tissues, and H.L. performed cellular experiments. H.L. and E.H.-Y.C. performed RNA extractions, and H.C. performed alignment of raw sequencing data; H.L. performed RNA-seq analysis. H.C. and Z.L. supervised RNA-seq processing and analysis. H.L., E.H.-Y.C., and K.X. performed mouse genotyping, colony management, and survival and weight tracking. M.A.D. designed and generated the hCIC-L plasmid. H.L. and M.A.D. generated mutant constructs and performed additional cloning, with technical assistance from E.H.-Y.C. R.R. performed size-exclusion chromatography. S.V. performed mouse behavior experiments. B.S. processed lung tissue for histopathology and performed morphometric analysis. H.L. wrote the manuscript and prepared the figures with input from all authors. H.Y.Z. reviewed all data and edited the manuscript.

