## Supplemental data for "Functional divergence of Capicua isoforms explains differential tissue vulnerability in neurological disease"

**Supplementary Tables**

**Supplemental Table S1. Overlapping differentially expressed genes between *Cic-S*-KO and *Atxn1*-KO cortex datasets. Related to Supplemental Figure S7.**

**Supplemental Table S2. Overlapping differentially expressed genes between *Cic-L*-KO and *Atxn1*-KO cortex datasets. Related to Supplemental Figure S7.**

**Supplemental Table S3. Overlapping differentially expressed genes between *Cic-S*-KO and *Atxn1*-KO cerebellum datasets. Related to Supplemental Figure S9.**

**Supplemental Table S4. Overlapping differentially expressed genes between *Cic-L*-KO and *Atxn1*-KO cerebellum datasets. Related to Supplemental Figure S9.**

### Supplementary Figures

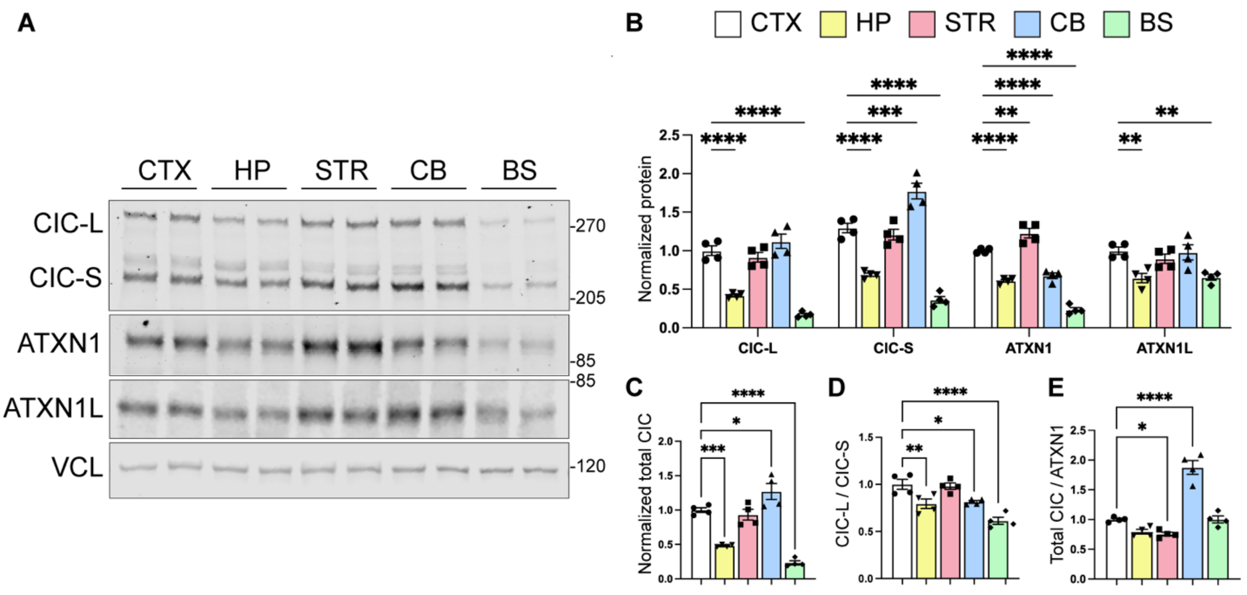


**Supplemental Figure S1. CIC-L, CIC-S, ATXN1, and ATXN1L expression in different brain regions.**

(A) Representative immunoblot of CIC-L, CIC-S, ATXN1, and ATXN1L in 12-week-old WT mouse brain lysates. CTX, cortex; HP, hippocampus; STR, striatum; CB, cerebellum; BS, brainstem.

(B, C) Quantification of each protein's expression: CIC-L, CIC-S, ATXN1, and ATXN1L (B); Total CIC combining CIC-L and CIC-S (C). Protein levels were normalized to Vinculin (VCL), and values were normalized to the cortex value for each respective protein.

(D) Ratio of CIC-L to CIC-S at each region.

(E) Total CIC was normalized to ATXN1 for each region. Mean of each region compared to the normalized cortical expression of each protein. Data are mean ± SEM; n = 4 per region. One-way ANOVA with Dunnett's multiple comparisons for (B-E); (*) $p < 0.05$, (**) $p < 0.01$, (***) $p < 0.001$, (****) $p < 0.0001$.

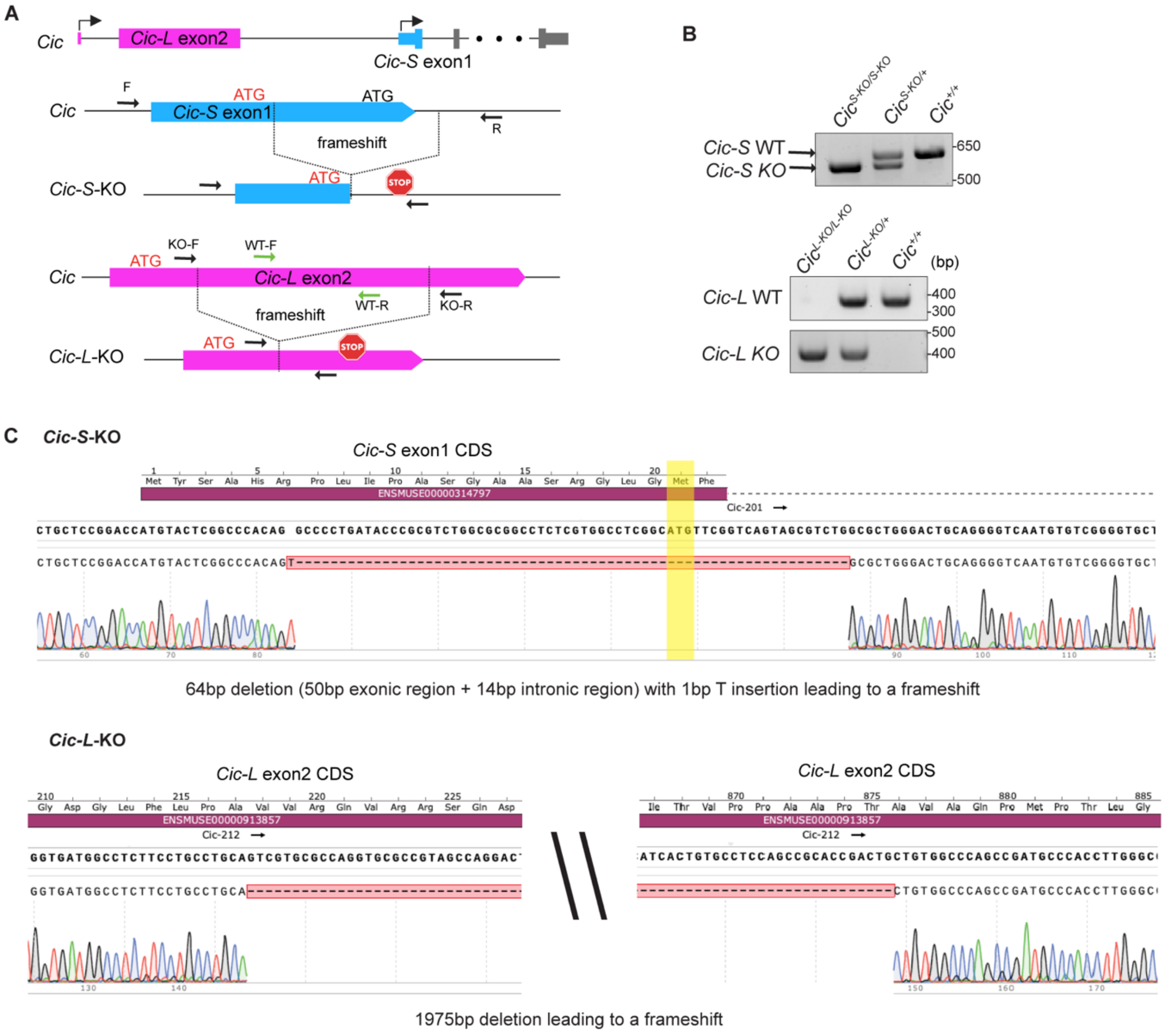
A
Cic
Cic-L exon2
Cic-S exon1
F
ATG
Cic-S exon1
ATG
R
frameshift
Cic-S-KO
ATG
STOP
KO-F
WT-F
ATG
Cic-L exon2
WT-R
KO-R
frameshift
Cic-L-KO
ATG
STOP
B
Cic S-KO/S-KO
Cic S-KO/+
Cic +/+
Cic-S WT
Cic-S KO
650
500
Cic L-KO/L-KO
Cic L-KO/+
Cic +/+
(bp)
Cic-L WT
400
300
Cic-L KO
500
400
C
Cic-S-KO
Cic-S exon1 CDS
ENSMUSE00000314797
Cic-201
64bp deletion (50bp exonic region + 14bp intronic region) with 1bp T insertion leading to a frameshift
Cic-L-KO
Cic-L exon2 CDS
ENSMUSE00000913857
Cic-212
Cic-L exon2 CDS
ENSMUSE00000913857
Cic-212
1975bp deletion leading to a frameshift

**Supplemental Figure S2. *Cic-S*-KO and *Cic-L*-KO were generated by frameshift indels.**

(A) Schematic of *Cic-L*-KO and *Cic-S*-KO allele generation. *Cic-L*-KO was generated by a 1975bp deletion of the first coding exon of *Cic-L* (*Cic-L* exon2). The deletion is downstream of the start codon and leads to a frameshift of *Cic-L*. *Cic-S*-KO was generated by an indel that spans the *Cic-S* unique exon1 and the downstream intron. The deletion is downstream of the start codon and spans 50bp of the exon including the downstream start codon and 14bp of the intron with a one bp insertion. Overall, this leads to a frameshift of *Cic-S*. The start codon is marked with red and the downstream ATG is marked with black. The PCR primers for genotyping are marked as arrows.

(B) Representative gel image of genotyping PCR reaction. *Cic-L* KO allele (383bp), *Cic-L* KO allele (403bp), *Cic-S* WT allele (633bp), and *Cic-S* KO allele (570bp).

(C) Sanger sequencing confirming a frameshift indel of *Cic-S*-KO (top), and *Cic-L*-KO (bottom) genomic DNA. The downstream ATG in *Cic-S* exon 1 is highlighted in yellow.

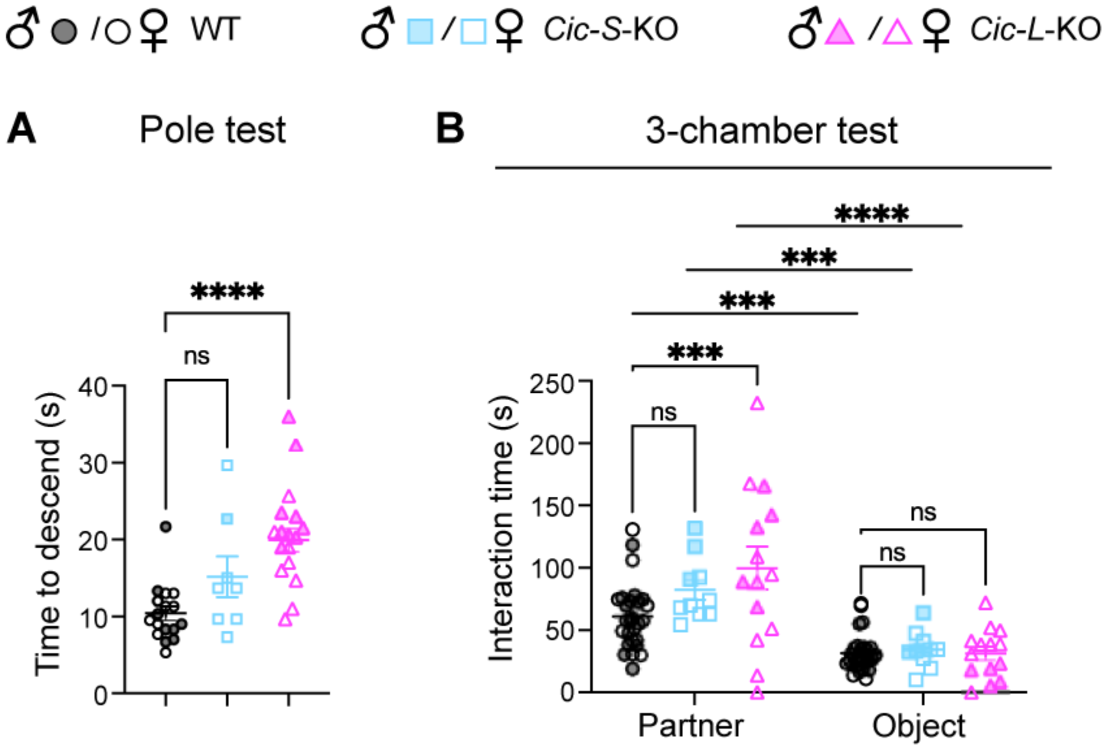


**Supplemental Figure S3. *Cic-L*-KO shows motor deficits in the pole test and more time spent with partner during a social interaction assay.**

(A) Fine motor behavior measured with pole test at 15-17 weeks of age.

(B) Social behavior measured with the three-chamber assay at 12-15 weeks of age.

For (A) n = 17 (8 males, 9 females) WT, n = 8 *Cic-S*-KO (1 males, 7 females), and n =16 *Cic-L*-KO (9 males, 7 females) were analyzed with one-way ANOVA with Tukey's multiple comparisons. For (B) n = 27 (12 males, 15 females) WT, n = 10 *Cic-S*-KO (3 males, 7 females), and n =14 *Cic-L*-KO (5 males, 9 females). Data are mean ± SEM. One-way ANOVA with Dunnett's multiple comparisons for (A); Two-way ANOVA with Dunnett's multiple comparisons for (B); (***) $p < 0.001$, (****) $p < 0.0001$, (ns) $p > 0.05$.

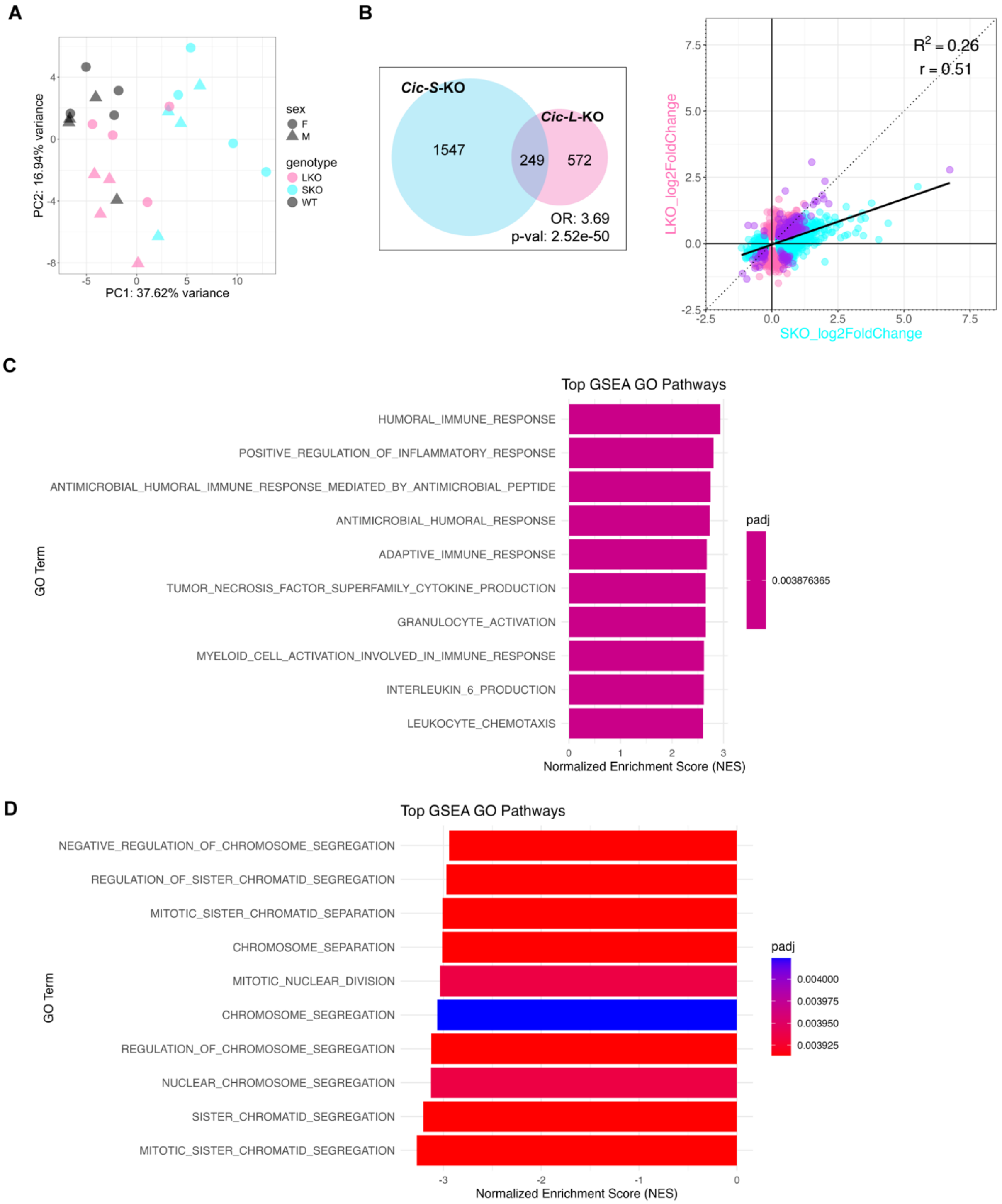

A
B
C
D
PC1: 37.62% variance
PC2: 16.94% variance
sex
F
M
genotype
LKO
SKO
WT
Cic-S-KO
Cic-L-KO
1547
249
572
OR: 3.69
p-val: 2.52e-50
R2 = 0.26
r = 0.51
SKO_log2FoldChange
LKO_log2FoldChange
Top GSEA GO Pathways
HUMORAL_IMMUNE_RESPONSE
POSITIVE_REGULATION_OF_INFLAMMATORY_RESPONSE
ANTIMICROBIAL_HUMORAL_IMMUNE_RESPONSE_MEDIATED_BY_ANTIMICROBIAL_PEPTIDE
ANTIMICROBIAL_HUMORAL_RESPONSE
ADAPTIVE_IMMUNE_RESPONSE
TUMOR_NECROSIS_FACTOR_SUPERFAMILY_CYTOKINE_PRODUCTION
GRANULOCYTE_ACTIVATION
MYELOID_CELL_ACTIVATION_INVOLVED_IN_IMMUNE_RESPONSE
INTERLEUKIN_6_PRODUCTION
LEUKOCYTE_CHEMOTAXIS
GO Term
padj
0.003876365
Normalized Enrichment Score (NES)
Top GSEA GO Pathways
NEGATIVE_REGULATION_OF_CHROMOSOME_SEGREGATION
REGULATION_OF_SISTER_CHROMATID_SEGREGATION
MITOTIC_SISTER_CHROMATID_SEPARATION
CHROMOSOME_SEPARATION
MITOTIC_NUCLEAR_DIVISION
CHROMOSOME_SEGREGATION
REGULATION_OF_CHROMOSOME_SEGREGATION
NUCLEAR_CHROMOSOME_SEGREGATION
SISTER_CHROMATID_SEGREGATION
MITOTIC_SISTER_CHROMATID_SEGREGATION
GO Term
padj
0.004000
0.003975
0.003950
0.003925
Normalized Enrichment Score (NES)

**Supplemental Figure S4. *Cic-S*-KO shows greater number and magnitude of DEGs in the lungs compared to *Cic-L*-KO DEGs.**

(A) Principal component analysis (PCA) of RNA-seq data from P6 lungs (WT, *Cic-S*-KO [SKO], *Cic-L*-KO [LKO]) using the top 500 most variable genes.

(B) Left: Venn diagram showing overlap of differentially expressed genes (DEGs; padj < 0.05 & |log2 fold change (log2FC)| >0.25) between *Cic-S*-KO and *Cic-L*-KO. Odds ratio (OR) and Fisher's exact *p*-value are indicated. A total of 15473 genes were analyzed. Right: Scatterplot comparing log2FC of DEGs *Cic-S*-KO versus *Cic-L*-KO. Pearson correlation coefficient (r) and $R^2$ are shown. Overlapping DEGs marked in purple.

(C, D) Top gene ontology (GO) enrichment terms for biological processes of *Cic-S*-KO (C) and *Cic-L*-KO (D).

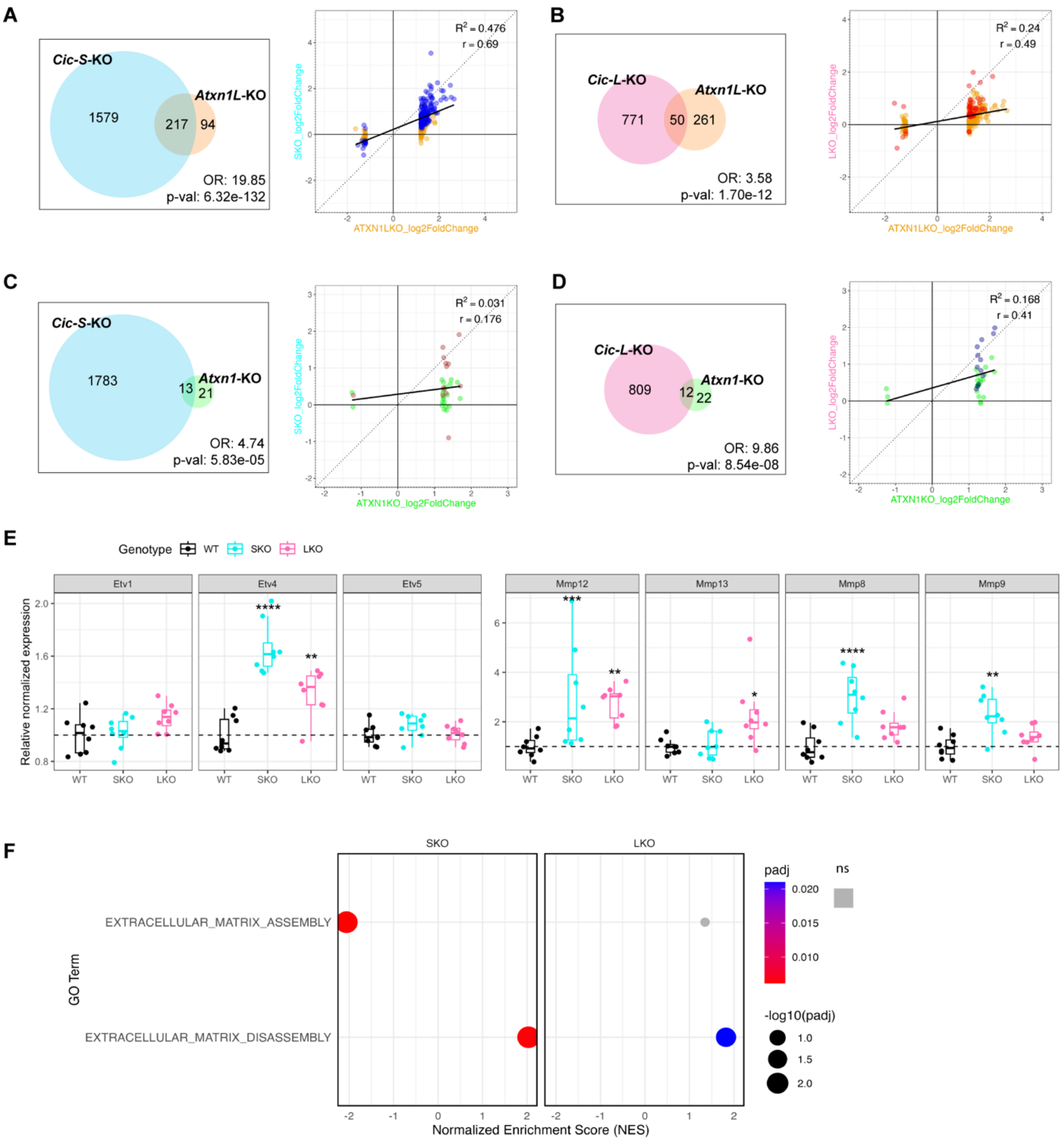
A
Cic-S-KO
Atxn1L-KO
1579
217
94
OR: 19.85
p-val: 6.32e-132
R2 = 0.476
r = 0.69
SKO_log2FoldChange
ATXN1LKO_log2FoldChange
B
Cic-L-KO
Atxn1L-KO
771
50
261
OR: 3.58
p-val: 1.70e-12
R2 = 0.24
r = 0.49
LKO_log2FoldChange
ATXN1LKO_log2FoldChange
C
Cic-S-KO
Atxn1-KO
1783
13
21
OR: 4.74
p-val: 5.83e-05
R2 = 0.031
r = 0.176
SKO_log2FoldChange
ATXN1KO_log2FoldChange
D
Cic-L-KO
Atxn1-KO
809
12
22
OR: 9.86
p-val: 8.54e-08
R2 = 0.168
r = 0.41
LKO_log2FoldChange
ATXN1KO_log2FoldChange
E
Genotype WT SKO LKO
Etv1
Etv4
Etv5
Mmp12
Mmp13
Mmp8
Mmp9
Relative normalized expression
WT SKO LKO
F
SKO
LKO
GO Term
EXTRACELLULAR_MATRIX_ASSEMBLY
EXTRACELLULAR_MATRIX_DISASSEMBLY
Normalized Enrichment Score (NES)
padj
0.020
0.015
0.010
ns
-log10(padj)
1.0
1.5
2.0

**Supplemental Figure S5. *Cic-S*-KO DEGs show greater overlap with *Atxn1l*-KO DEGs than *Cic-L*-KO DEGs in lung.**

(A, B) Left: Venn diagrams showing overlap between DEGs in *Cic-S*-KO [SKO] (A) or *Cic-L*-KO [LKO] (B) and DEGs from *Atxn1l*-KO lungs reported in *Lee et al., 2011*. Odds ratio (OR) and Fisher's exact test p-value are indicated. A total of 15473 genes were analyzed. Right: Scatterplot comparing log2FC of DEGs of *Cic-S*-KO (A) or *Cic-L*-KO (B) versus *Atxn1l*-KO. Pearson correlation coefficient (r) and $R^2$ are shown. Overlapping DEGs marked in blue (A) or red (B).

(C, D) Left: Venn diagrams showing overlap between DEGs in *Cic-S*-KO (A) or *Cic-L*-KO (B) and DEGs from *Atxn1*-KO lungs reported in *Lee et al., 2011*. OR and Fisher's exact *p*-value are indicated. A total of 15473 genes were analyzed. Right: Scatterplot comparing log2FC of DEGs of *Cic-S*-KO (C) or *Cic-L*-KO (D) versus *Atxn1*-KO. Pearson correlation coefficient (r) and $R^2$ are shown. Overlapping DEGs marked in brown (C) or navy (D).

(E) Boxplot of *Etv-Mmp* genes. Differential expression statistics were obtained using DESeq2; (*) adjusted $p < 0.05$, (**) adjusted $p < 0.01$, (***), adjusted $p < 0.001$, (****) adjusted $p < 0.0001$.

(F) Enrichment of GO of extracellular matrix assembly and disassembly terms of *Cic-S*-KO (Left) or *Cic-L*-KO (Right) gene set enrichment analysis (GSEA).

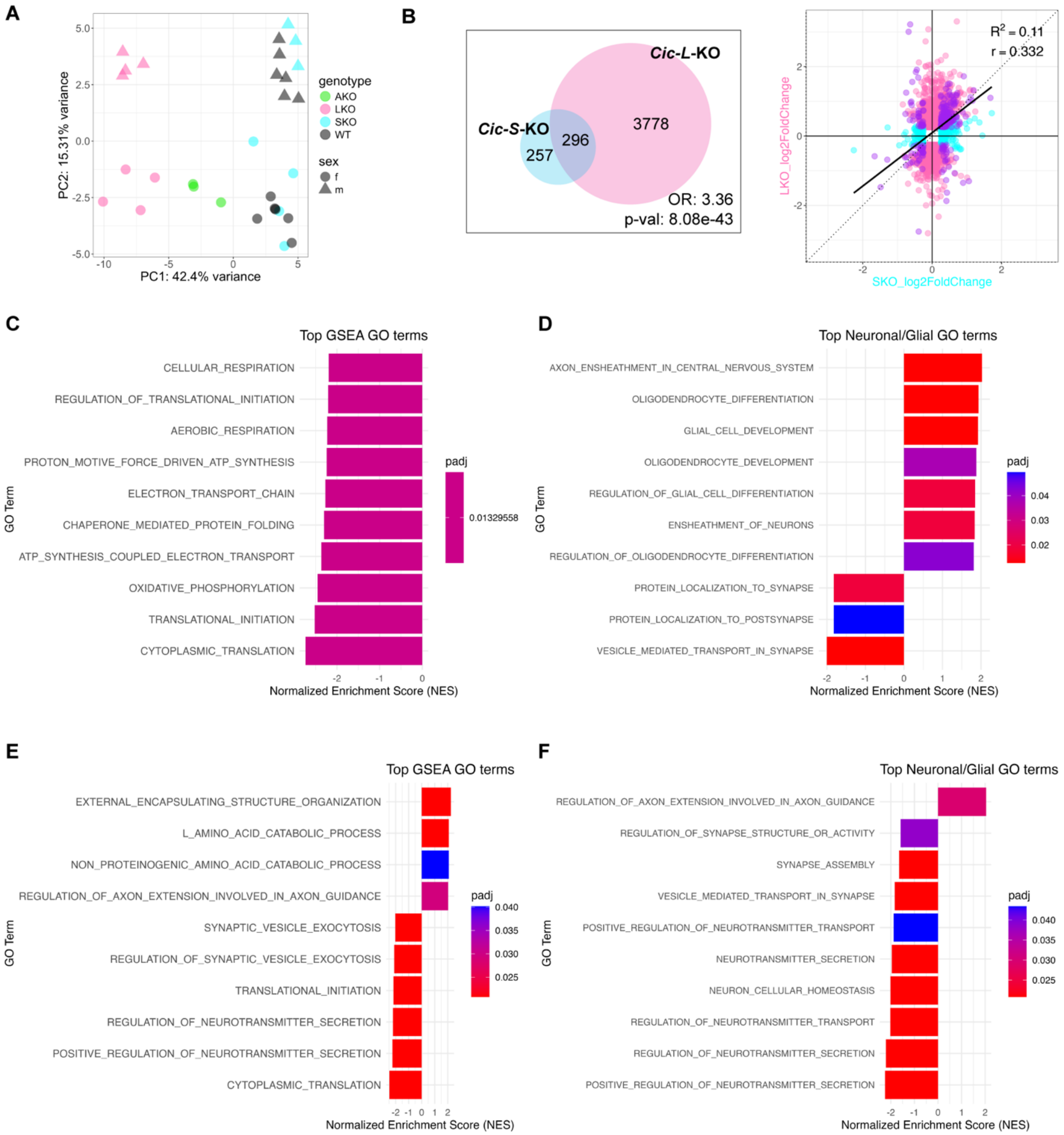
A
B
C
D
E
F
genotype
AKO
LKO
SKO
WT
sex
f
m
PC1: 42.4% variance
PC2: 15.31% variance
Cic-S-KO
Cic-L-KO
257
296
3778
OR: 3.36
p-val: 8.08e-43
R2 = 0.11
r = 0.332
SKO_log2FoldChange
LKO_log2FoldChange
Top GSEA GO terms
CELLULAR_RESPIRATION
REGULATION_OF_TRANSLATIONAL_INITIATION
AEROBIC_RESPIRATION
PROTON_MOTIVE_FORCE_DRIVEN_ATP_SYNTHESIS
ELECTRON_TRANSPORT_CHAIN
CHAPERONE_MEDIATED_PROTEIN_FOLDING
ATP_SYNTHESIS_COUPLED_ELECTRON_TRANSPORT
OXIDATIVE_PHOSPHORYLATION
TRANSLATIONAL_INITIATION
CYTOPLASMIC_TRANSLATION
padj
0.01329558
GO Term
Normalized Enrichment Score (NES)
Top Neuronal/Glial GO terms
AXON_ENSHEATHMENT_IN_CENTRAL_NERVOUS_SYSTEM
OLIGODENDROCYTE_DIFFERENTIATION
GLIAL_CELL_DEVELOPMENT
OLIGODENDROCYTE_DEVELOPMENT
REGULATION_OF_GLIAL_CELL_DIFFERENTIATION
ENSHEATHMENT_OF_NEURONS
REGULATION_OF_OLIGODENDROCYTE_DIFFERENTIATION
PROTEIN_LOCALIZATION_TO_SYNAPSE
PROTEIN_LOCALIZATION_TO_POSTSYNAPSE
VESICLE_MEDIATED_TRANSPORT_IN_SYNAPSE
Top GSEA GO terms
EXTERNAL_ENCAPSULATING_STRUCTURE_ORGANIZATION
L_AMINO_ACID_CATABOLIC_PROCESS
NON_PROTEINOGENIC_AMINO_ACID_CATABOLIC_PROCESS
REGULATION_OF_AXON_EXTENSION_INVOLVED_IN_AXON_GUIDANCE
SYNAPTIC_VESICLE_EXOCYTOSIS
REGULATION_OF_SYNAPTIC_VESICLE_EXOCYTOSIS
TRANSLATIONAL_INITIATION
REGULATION_OF_NEUROTRANSMITTER_SECRETION
POSITIVE_REGULATION_OF_NEUROTRANSMITTER_SECRETION
CYTOPLASMIC_TRANSLATION
Top Neuronal/Glial GO terms
REGULATION_OF_AXON_EXTENSION_INVOLVED_IN_AXON_GUIDANCE
REGULATION_OF_SYNAPSE_STRUCTURE_OR_ACTIVITY
SYNAPSE_ASSEMBLY
VESICLE_MEDIATED_TRANSPORT_IN_SYNAPSE
POSITIVE_REGULATION_OF_NEUROTRANSMITTER_TRANSPORT
NEUROTRANSMITTER_SECRETION
NEURON_CELLULAR_HOMEOSTASIS
REGULATION_OF_NEUROTRANSMITTER_TRANSPORT
REGULATION_OF_NEUROTRANSMITTER_SECRETION
POSITIVE_REGULATION_OF_NEUROTRANSMITTER_SECRETION

**Supplemental Figure S6.** ***Cic-L*****-KO shows greater number and magnitude of DEGs than** ***Cic-S*****-KO in the cortex.**

(A) Principal component analysis (PCA) of RNA-seq data from 12-week-old cortex of WT, *Cic-S*-KO [SKO], *Cic-L*-KO [LKO] of this study and 10-week-old cortex from *(unpublished data)*. *Atxn1*-KO [AKO] and WT using the top 500 most variable genes.

(B) Left: Venn diagram showing overlap of DEGs between *Cic-S*-KO and *Cic-L*-KO. OR and Fisher's exact *p*-value are indicated. A total of 15354 genes were analyzed. Right: Scatterplot comparing log2FC of DEGs *Cic-S*-KO versus *Cic-L*-KO. Pearson correlation coefficient (r) and $R^2$ are shown. Overlapping DEGs marked in purple.

(C, D) Top gene ontology (GO) terms of all biological processes (C) and neuro- or glia-related terms (D) of *Cic-S*-KO gene set enrichment analysis (GSEA).

(E, F) Top enriched GO terms of all biological process (E) and neuro- or glia- related -terms (F) of *Cic-L*-KO GSEA.

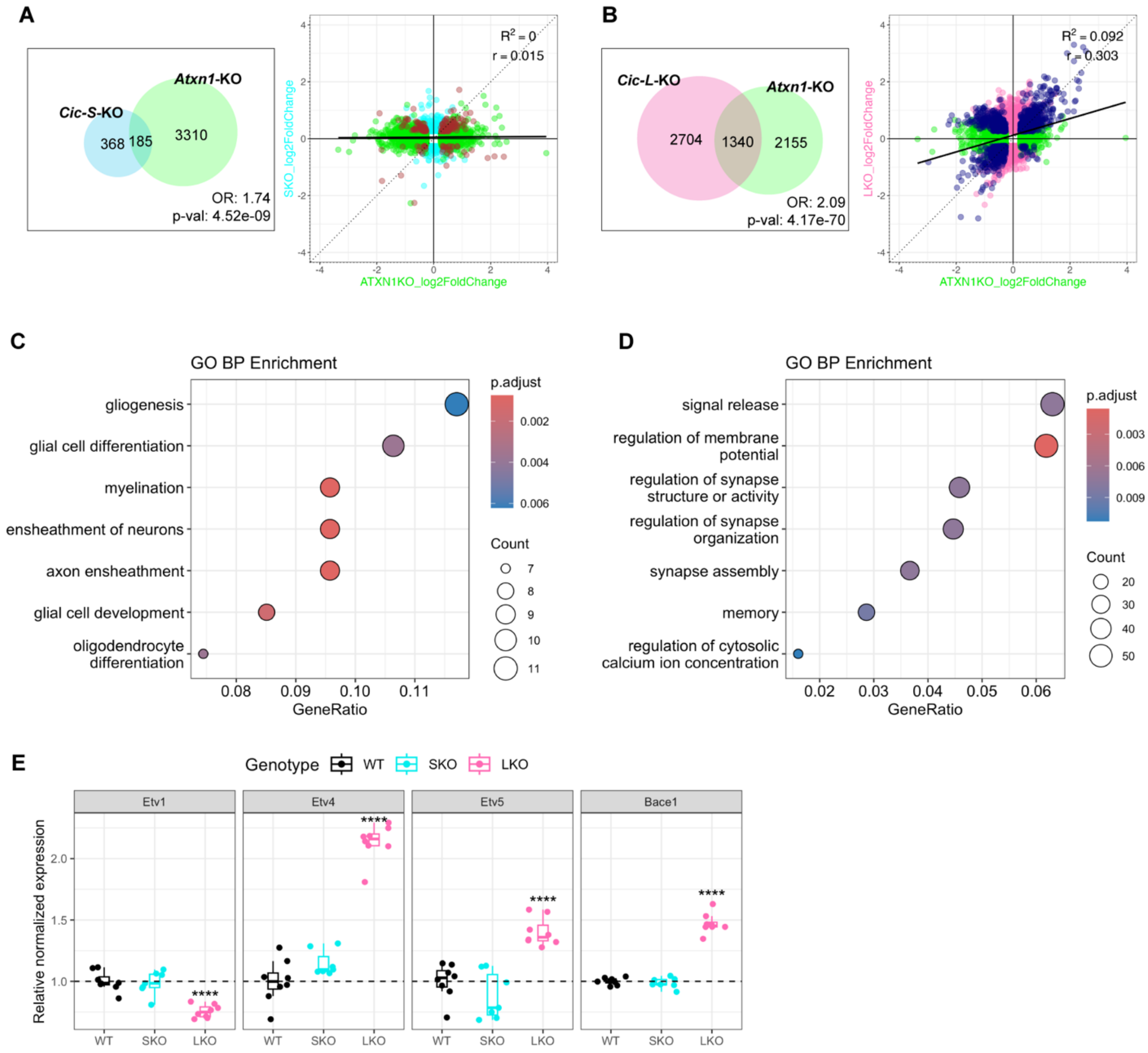
A
Cic-S-KO
Atxn1-KO
368
185
3310
OR: 1.74
p-val: 4.52e-09
$R^2 = 0$
r = 0.015
SKO_log2FoldChange
ATXN1KO_log2FoldChange
B
Cic-L-KO
Atxn1-KO
2704
1340
2155
OR: 2.09
p-val: 4.17e-70
$R^2 = 0.092$
r = 0.303
LKO_log2FoldChange
ATXN1KO_log2FoldChange
C
GO BP Enrichment
gliogenesis
glial cell differentiation
myelination
ensheathment of neurons
axon ensheathment
glial cell development
oligodendrocyte differentiation
GeneRatio
p.adjust
Count
D
GO BP Enrichment
signal release
regulation of membrane potential
regulation of synapse structure or activity
regulation of synapse organization
synapse assembly
memory
regulation of cytosolic calcium ion concentration
GeneRatio
p.adjust
Count
E
Genotype WT SKO LKO
Etv1
Etv4
Etv5
Bace1
Relative normalized expression
WT SKO LKO
****

**Supplemental Figure S7. *Cic-L*-KO DEGs show greater overlap with *Atxn1*-KO DEGs than *Cic-S*-KO DEGs in the cortex.**

(A, B) Left: Venn diagrams showing overlap between DEGs in *Cic-S*-KO [SKO] (A) or *Cic-L*-KO [LKO] (B) and DEGs (padj < 0.05 & |log2FC| >0.25) from *Atxn1*-KO cortex (*unpublished data)*. OR and Fisher's exact *p*-value are indicated. A total of 15290 genes were analyzed. Right: Scatterplot comparing log2FC of DEGs of *Cic-S*-KO (A) or *Cic-L*-KO (B) versus *Atxn1*-KO. Pearson correlation coefficient (r) and $R^2$ are shown. Overlapping DEGs marked in brown (A) or navy (B).

(C, D) Top enriched gene ontology terms of biological process of the DEGs that overlap and change in the same direction between *Atxn1*-KO and *Cic-S*-KO (C), or *Atxn1*-KO and *Cic-L*-KO (D).

(E) Boxplot of *Etv1/4/5-Bace1*. Differential expression statistics were obtained using DESeq2; (****) adjusted p < 0.0001.

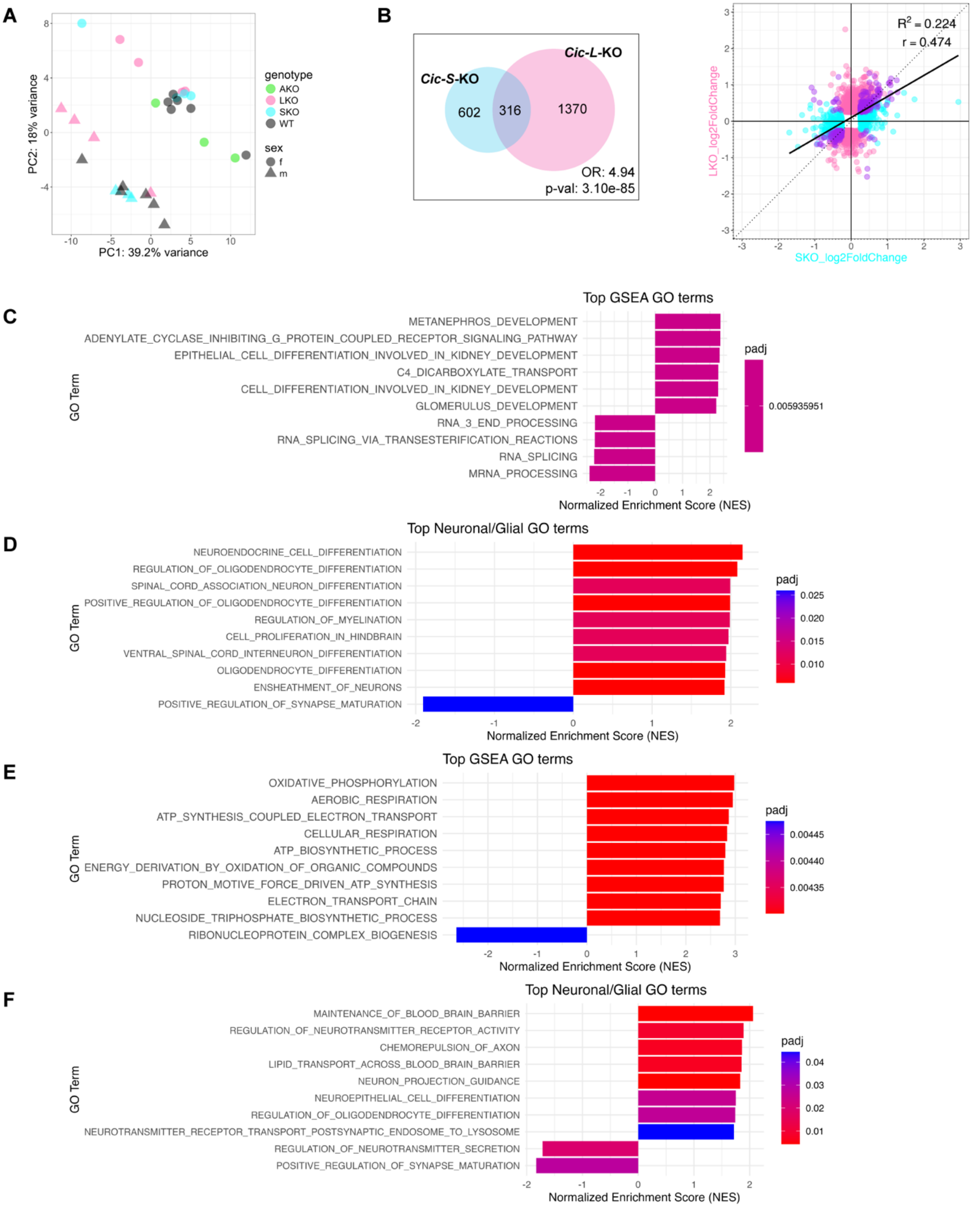

A
PC1: 39.2% variance
PC2: 18% variance
genotype
AKO
LKO
SKO
WT
sex
f
m
B
Cic-S-KO
Cic-L-KO
602
316
1370
OR: 4.94
p-val: 3.10e-85
$R^2$ = 0.224
r = 0.474
LKO_log2FoldChange
SKO_log2FoldChange
C
Top GSEA GO terms
METANEPHROS_DEVELOPMENT
ADENYLATE_CYCLASE_INHIBITING_G_PROTEIN_COUPLED_RECEPTOR_SIGNALING_PATHWAY
EPITHELIAL_CELL_DIFFERENTIATION_INVOLVED_IN_KIDNEY_DEVELOPMENT
C4_DICARBOXYLATE_TRANSPORT
CELL_DIFFERENTIATION_INVOLVED_IN_KIDNEY_DEVELOPMENT
GLOMERULUS_DEVELOPMENT
RNA_3_END_PROCESSING
RNA_SPLICING_VIA_TRANSESTERIFICATION_REACTIONS
RNA_SPLICING
MRNA_PROCESSING
GO Term
Normalized Enrichment Score (NES)
padj
0.005935951
D
Top Neuronal/Glial GO terms
NEUROENDOCRINE_CELL_DIFFERENTIATION
REGULATION_OF_OLIGODENDROCYTE_DIFFERENTIATION
SPINAL_CORD_ASSOCIATION_NEURON_DIFFERENTIATION
POSITIVE_REGULATION_OF_OLIGODENDROCYTE_DIFFERENTIATION
REGULATION_OF_MYELINATION
CELL_PROLIFERATION_IN_HINDBRAIN
VENTRAL_SPINAL_CORD_INTERNEURON_DIFFERENTIATION
OLIGODENDROCYTE_DIFFERENTIATION
ENSHEATHMENT_OF_NEURONS
POSITIVE_REGULATION_OF_SYNAPSE_MATURATION
GO Term
Normalized Enrichment Score (NES)
padj
0.025
0.020
0.015
0.010
E
Top GSEA GO terms
OXIDATIVE_PHOSPHORYLATION
AEROBIC_RESPIRATION
ATP_SYNTHESIS_COUPLED_ELECTRON_TRANSPORT
CELLULAR_RESPIRATION
ATP_BIOSYNTHETIC_PROCESS
ENERGY_DERIVATION_BY_OXIDATION_OF_ORGANIC_COMPOUNDS
PROTON_MOTIVE_FORCE_DRIVEN_ATP_SYNTHESIS
ELECTRON_TRANSPORT_CHAIN
NUCLEOSIDE_TRIPHOSPHATE_BIOSYNTHETIC_PROCESS
RIBONUCLEOPROTEIN_COMPLEX_BIOGENESIS
GO Term
Normalized Enrichment Score (NES)
padj
0.00445
0.00440
0.00435
F
Top Neuronal/Glial GO terms
MAINTENANCE_OF_BLOOD_BRAIN_BARRIER
REGULATION_OF_NEUROTRANSMITTER_RECEPTOR_ACTIVITY
CHEMOREPULSION_OF_AXON
LIPID_TRANSPORT_ACROSS_BLOOD_BRAIN_BARRIER
NEURON_PROJECTION_GUIDANCE
NEUROEPITHELIAL_CELL_DIFFERENTIATION
REGULATION_OF_OLIGODENDROCYTE_DIFFERENTIATION
NEUROTRANSMITTER_RECEPTOR_TRANSPORT_POSTSYNAPTIC_ENDOSOME_TO_LYSOSOME
REGULATION_OF_NEUROTRANSMITTER_SECRETION
POSITIVE_REGULATION_OF_SYNAPSE_MATURATION
GO Term
Normalized Enrichment Score (NES)
padj
0.04
0.03
0.02
0.01

**Supplemental Figure S8. *Cic-L*-KO shows greater number and magnitude of DEGs in the cerebellum than *Cic-S*-KO.**

(A) Principal component analysis (PCA) of RNA-seq data from 12-week-old cerebellum of WT, *Cic-S*-KO [SKO], *Cic-L*-KO [LKO] of this study and 10-week-old cerebellum *Atxn1*-KO [AKO] and WT (*unpublished data*) using the top 500 most variable genes.

(B) Left: Venn diagram showing overlap of DEGs between *Cic-S*-KO and *Cic-L*-KO. OR and Fisher's exact *p*-value are indicated. A total of 15354 genes were analyzed. Right: Scatterplot comparing log2FC of DEGs *Cic-S*-KO versus *Cic-L*-KO. Pearson correlation coefficient (r) and $R^2$ are shown. Overlapping DEGs marked in purple.

(C, D) Top gene ontology (GO) terms of all biological process (C) and neuro- or glia- related terms (D) of *Cic-S*-KO gene set enrichment analysis (GSEA).

(E, F) Top enriched GO terms of all biological process (E) and neuro- or glia- related -terms (F) of *Cic-L*-KO GSEA.

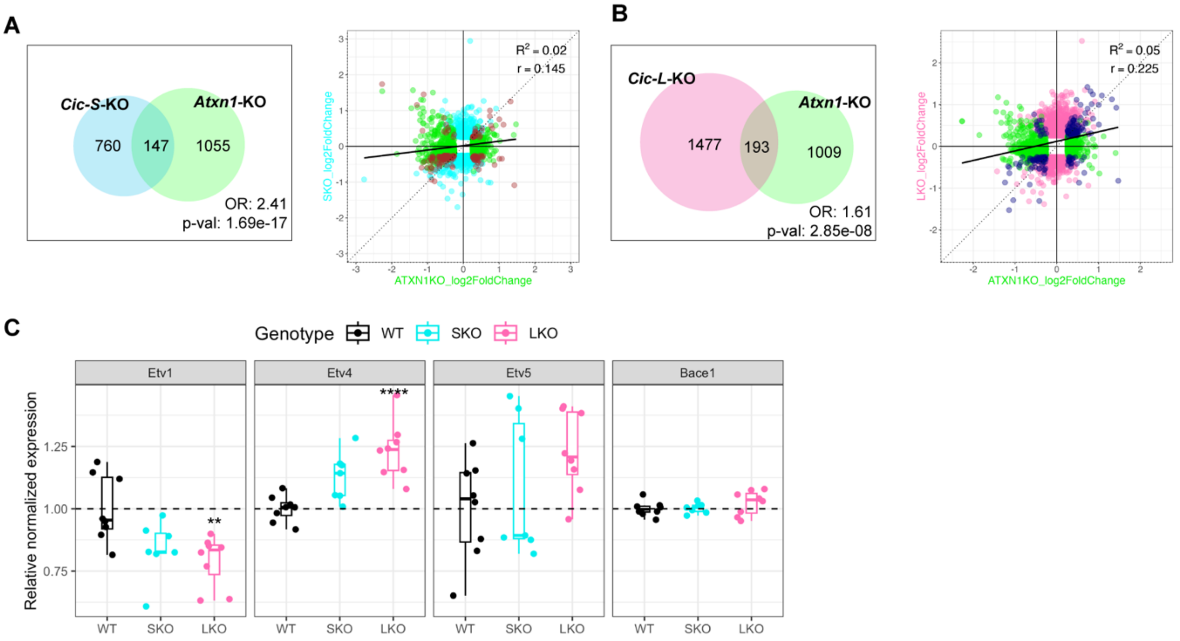


**Supplemental Figure S9. *Cic-S*-KO DEGs show greater overlap in the cerebellum with *Atxn1*-KO DEGs than *Cic-L*-KO DEGs.**

(A, B) Left: Venn diagrams showing overlap between DEGs in *Cic-S*-KO [SKO] (A) or *Cic-L*-KO [LKO] (B) and DEGs (padj < 0.05 & |log2FC| >0.25) from *Atxn1*-KO cerebellum (*unpublished data*). OR and Fisher's exact *p*-value are indicated. A total of 15290 genes were analyzed. Right: Scatterplot comparing log2FC of DEGs of *Cic-S*-KO (A) or *Cic-L*-KO (B) versus *Atxn1*-KO. Pearson correlation coefficient (r) and $R^2$ are shown. Overlapping DEGs marked in brown (A) or navy (B).

(C) Boxplot of *Etv1/4/5-Bace1*. Differential expression statistics were obtained using DESeq2; (**) adjusted p < 0.01, (****) adjusted p < 0.0001.

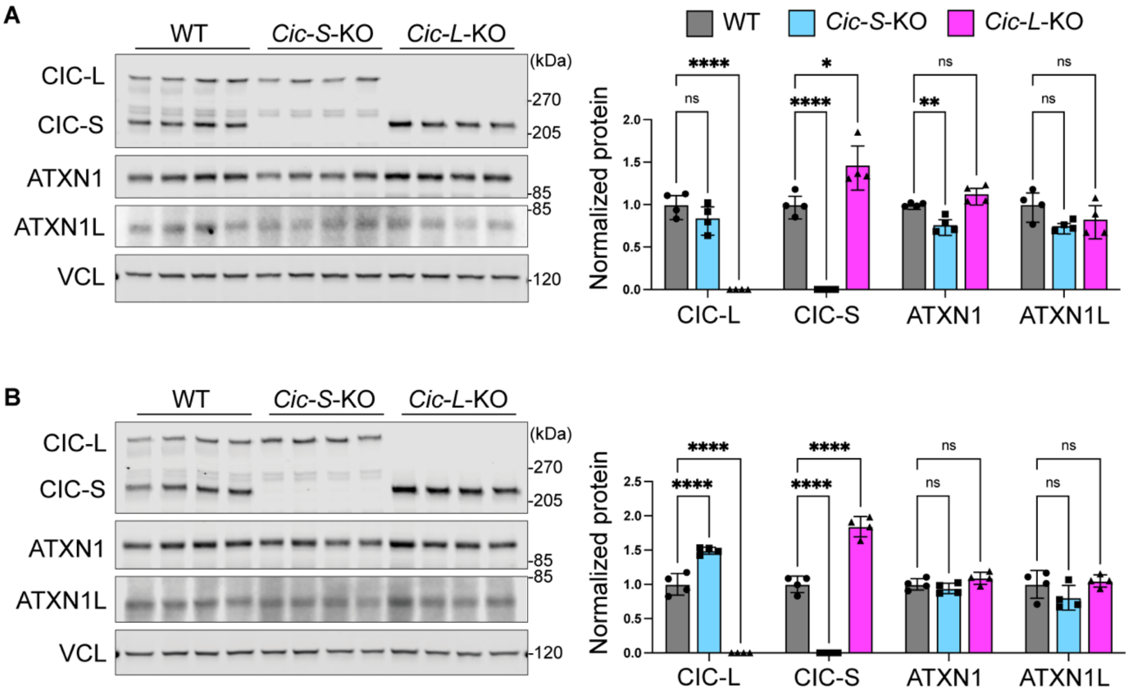


**Supplemental Figure S10. CIC, ATXN1, and ATXN1L expression in cortex and cerebellum of WT, *Cic-S*-KO, and *Cic-L*-KO mice.**

(A, B) Representative immunoblots and quantification of CIC-L, CIC-S, ATXN1, and ATXN1L protein levels in cortex (A) and cerebellum (B) of 12-week-old WT, *Cic-S*-KO, and *Cic-L*-KO mice. Protein levels were normalized to VCL, and values were normalized to the WT. Data are mean ± SEM; n = 4 per region. One-way ANOVA with Dunnett's multiple comparisons; (*) $p < 0.05$, (**) $p < 0.01$, (***) $p < 0.001$, (****) $p < 0.0001$, (ns) $p>0.05$.

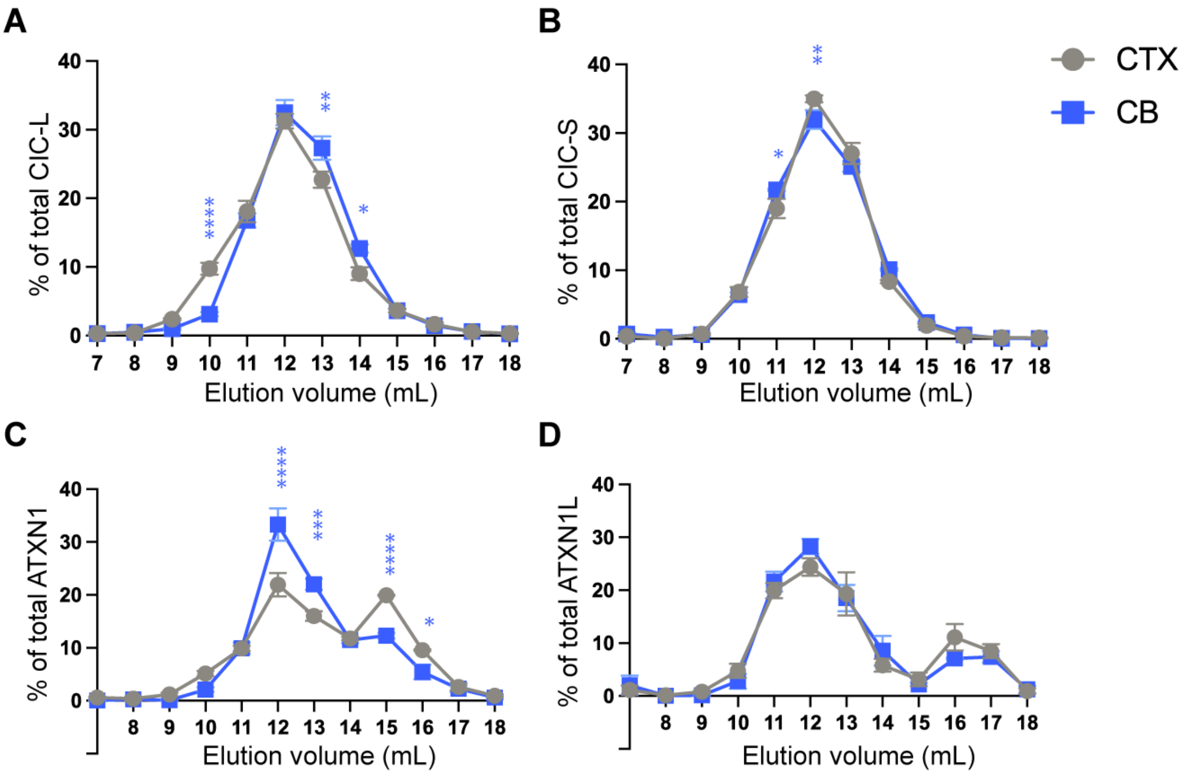


**Supplemental Figure S11. CIC isoforms form different complexes with ATXN1 and ATXN1L in the WT cortex and cerebellum (related to Figure 5).**

(A-D) SEC elution profile of each protein of WT cortex and cerebellum. The percentage of each protein (mean ± SEM) in each fraction was determined from n = 3–4 independent cortical or cerebellar extracts. Protein abundance in each fraction was quantified by densitometry, with the signal in each fraction normalized to the total signal across all fractions for both regions. Two-way ANOVA with Šídák’s multiple comparisons; (*) $p < 0.05$, (**) $p < 0.01$, (***) $p < 0.001$, (****) $p < 0.0001$.

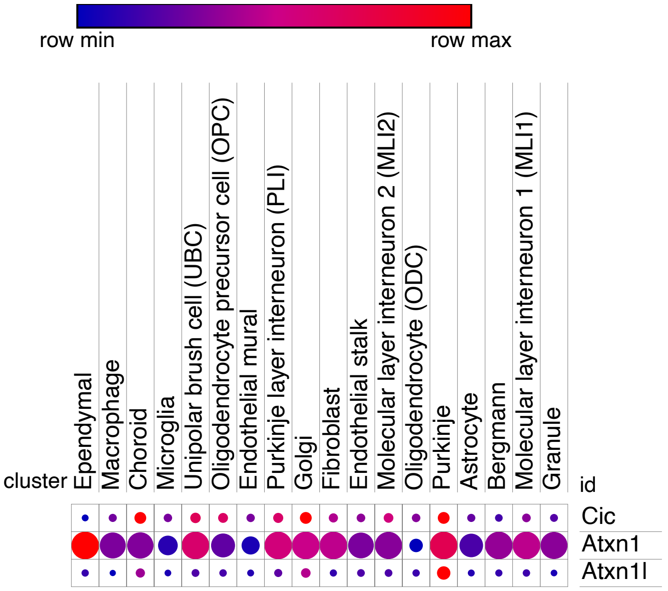


**Supplemental Figure S12.** ***Cic, Atxn1,*** **and** ***Atxn1l*** **expression in mouse cerebellum at P60.**

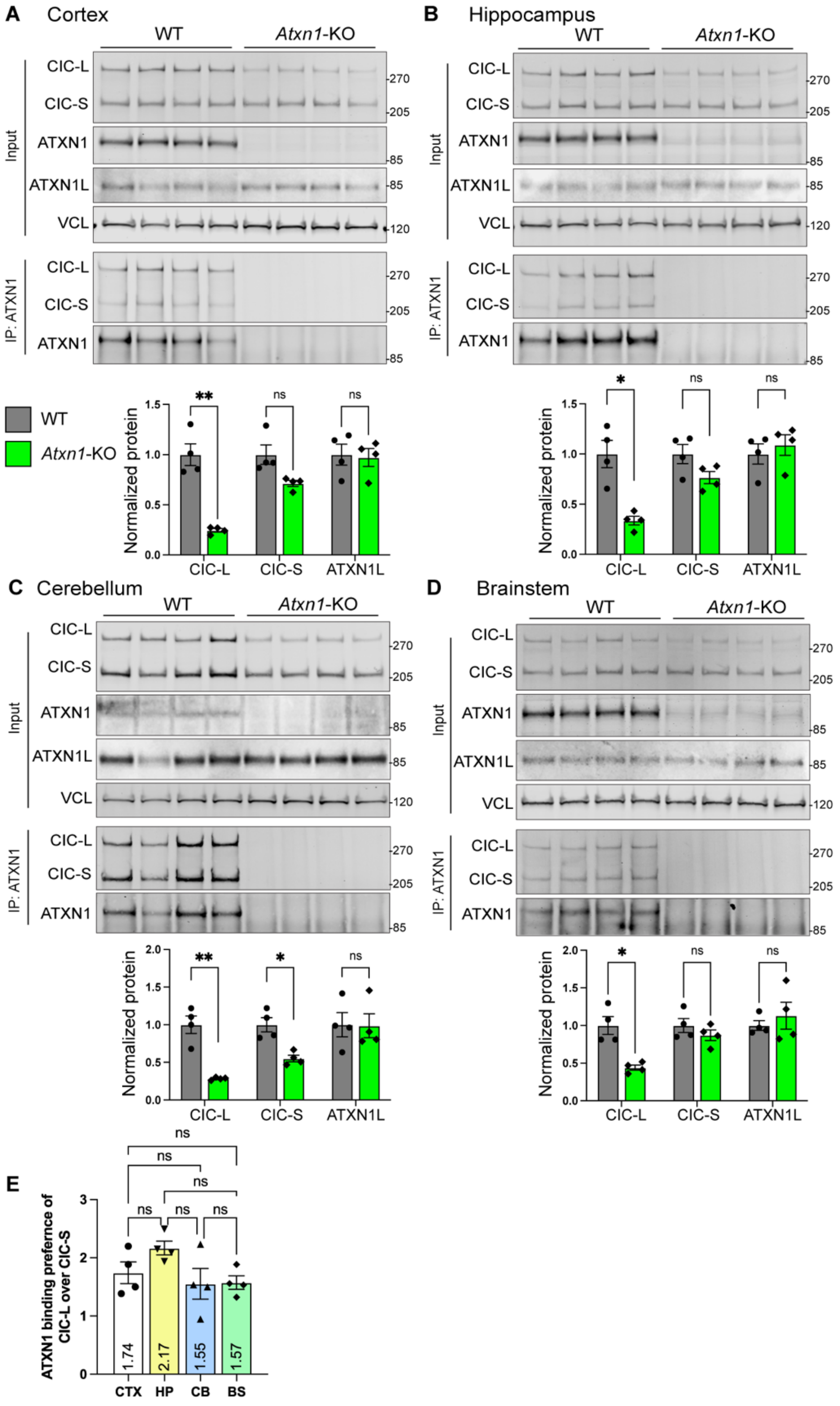

A Cortex
B Hippocampus
C Cerebellum
D Brainstem
WT
Atxn1-KO
Input
IP: ATXN1
CIC-L
CIC-S
ATXN1
ATXN1L
VCL
270
205
85
120
Normalized protein
**
*
ns
E
ATXN1 binding prefernce of CIC-L over CIC-S
1.74
2.17
1.55
1.57
CTX
HP
CB
BS

**Supplemental Figure S13. CIC-L is more dependent on ATXN1 than CIC-S in all brain regions.**

(A-D) Representative immunoblots (top) and quantification of input (bottom) of ATXN1, ATXN1L and CIC expression in WT mice and *Atxn1*-KO mice of four different brain regions. Protein levels were normalized to VCL, then relative levels were normalized to WT expression.

(E) Binding preference of CIC-L/CIC-S in ATXN1 IP normalized to the ratio in input. CTX, cortex; HP, hippocampus; CB, cerebellum; BS, brainstem. Data are mean ± SEM; $n = 4$ per region. Welch's t-test for (A-D); One-way ANOVA with Tukey's multiple comparisons for (E). (*) $p < 0.05$, (**) $p < 0.01$, (ns) $p > 0.05$.

### Supplementary Materials and Methods

#### Behavioral assays

**Pole test**: Mice (15-17 weeks of age) were placed head-upward at the top of a vertical threaded metal pole. Time to descend to the base was recorded with a 60 s cutoff. If a mouse turned but fell before reaching the floor, the time to fall was recorded. Mice unable to turn downward were scored as 60 s.

**Three-chamber sociability assay**: Partner mice (matched for age, sex, and genetic background) were habituated to the three-chamber apparatus (length: 54 cm; width: 40 cm; height: 23 cm) divided into three chambers by removable partitions, for 1 h per day on two consecutive days prior to testing, while placed inside wire cups. On the test day, following habituation, experimental mice (12-15 weeks of age) were placed in the central chamber and allowed to freely explore for 10 min. A novel partner mouse was then placed inside a wire cup in either the left or right chamber, while a novel object was placed inside a wire cup in the opposite chamber. Placement of the partner mouse was randomized across trials. Experimental mice were allowed to explore for 10 min, and interaction time with each wire cup was manually scored.

#### RNA extraction

Cortical and cerebellar tissues from 12-week-old mice, and lung tissues from P6 mice, were dissected, flash-frozen in liquid nitrogen, and stored at −80 °C until further processing. Total RNA was isolated using the miRNeasy Mini Kit (Qiagen, 217004) according to the manufacturer’s instructions. RNA concentration and purity were measured using a NanoDrop 1000 (Thermo Fisher).

**RNA sequencing and analysis**

RNA samples were submitted to Azenta Life Sciences for RNA integrity assessment, library preparation, and sequencing. Libraries were generated following polyA selection for mRNA and sequenced on Illumina NovaSeq or HiSeq platforms, yielding ~ 30 million 150-bp paired-end reads per sample.

For each sample, the raw reads from multiple lanes were merged, and then were aligned to the Mus musculus reference transcriptome (GRCm38.p6, GENCODE M18) using STAR v2.7 with "-quantMode GeneCounts" to generate raw read counts (Dobin et al. 2013).

Differential gene expression analysis was performed using DESeq2 v1.44.0 (Love et al. 2014). Principal component analysis (PCA) was performed on the top 500 most variable genes. Genes were considered differentially expressed if adjusted $p < 0.05$ and $|\log2 \text{ fold change}| > 0.25$.

Previously published lung P6 microarray data of *Atxn1*-KO and *Atxn1l*-KO were obtained from *Lee et al., 2011*. Differentially expressed genes (DEGs) reported in that dataset were used for comparison; the original publication defined DEGs using thresholds of $p < 0.05$ and $|\text{fold change}| > 1.2$. For *Atxn1*-KO RNA-seq data from cortex and cerebellum (*unpublished data*), read count data were processed with the same DESeq2 pipeline, and limma v3.60.6 (Ritchie et al. 2015) was used for batch correction prior to PCA.

Gene set enrichment analysis (GSEA) for Gene Ontology (GO) terms was conducted on all ranked genes using fgsea v1.30.0, with neuro/glia-related GO terms selected based on keywords including “neuron,” “neuro,” “nervous,” “synapse,” “axon,” “glia,” “glial,” “astrocyte,” “oligodendrocyte,” “microglia,” “myelin,” and “brain.” (Korotkevich et al. 2019) Overlapping differentially expressed genes were analyzed for GO term enrichment using clusterProfiler v4.12.6 (Xu et al. 2024), with the background universe defined as all genes detected in the RNA-seq dataset for the corresponding tissue.
